# Federated single-cell QTL meta-analysis reveals novel disease mechanisms

**DOI:** 10.64898/2026.01.20.700519

**Authors:** Daniel Kaptijn, Lieke Michielsen, Drew Neavin, Aida Ripoll-Cladellas, José Alquicira-Hernández, Maryna Korshevniuk, Jimmy Tsz Hang Lee, Roy Oelen, Martijn Vochteloo, Robert Warmerdam, Yoshinari Ando, Maria Ban, Odmaa Bayaraa, Marijn Berg, Irene van Blokland, Daniel Considine, Mame M. Dieng, Ryuya Edahiro, M. Grace Gordon, Hilde E. Groot, Pim van der Harst, Matthias Heinig, Chung-Chau Hon, Youssef Idaghdour, Pooja Kathail, Niek de Klein, Wenchao Li, Yang Li, Corinna Losert, Vinu Manikanda, Jonathan Moody, Haroon Naeem, Younes Mokrab, Martijn C. Nawijn, Mihai Netea, Jelmer Niewold, Yukinori Okada, Stephen Sawcer, Issiaka Soulama, Oliver Stegle, Yakov Tsepilov, Woong-Yang Park, Deepa Rajagopalan, Tala Shahin, Jay W. Shin, Gosia Trynka, Urmo Võsa, Harm-Jan Westra, Seyhan Yazar, Jimmie Ye, zhenhua zhang, eQTLGen consortium, Martin Hemberg, Ahmed Mahfouz, Marta Melé, Joseph E. Powell, Marc Jan Bonder, Lude Franke, Monique G.P. van der Wijst

## Abstract

Genetic effects on gene expression are often cell type-specific and obscured in bulk analyses. To resolve this context-dependent regulation, we performed a federated *cis-*eQTL meta-analysis across 12 PBMC datasets (2,032 individuals, 2.5 million cells). Across six immune cell types, we identified *cis-*eQTLs for 6,592 genes and fine-mapped 14,985 independent loci. Notably, the 42% of eQTLs that were undetected in a bulk eQTL study on 43,301 whole blood samples also showed stronger enrichment for disease GWAS loci. We further identified three genome-wide significant and 65 suggestive loci affecting the abundance of (rare) immune cell types and validated these using previously reported hematological GWAS and bulk-derived *trans-*eQTLs. Integrating single-cell *cis*-eQTLs with bulk *trans-*eQTLs enabled us to anchor 6,382 *trans-*eGenes (37.2% novel) to upstream regulators and reconstruct directed gene regulatory relationships. For example, a hemorrhoidal disease-associated variant showed a CD4+ T cell-specific *cis-*eQTL on *BACH1* that colocalized with 45 immune and metabolic *trans*-eGenes. These results demonstrate the power of single-cell QTL meta-analysis in interpreting complex trait genetics.

## Introduction

Genome-wide association studies (GWAS) have now revealed over 200,000 genetic risk factors, mostly single nucleotide polymorphisms (SNPs), for more than 200 different traits^1^. Deciphering how these genetic variants contribute to disease is fundamental for advancing therapeutic strategies. However, as the majority of these GWAS SNPs are in the non-coding part of the genome, it has been thus far challenging to pinpoint the molecular mechanisms underlying how these SNPs confer disease risk.

Linking GWAS SNPs to cell type abundance or specific molecular data layers through quantitative trait locus (QTL) analyses has already increased our understanding of these underlying mechanisms. Previously, the largest effort, involving 563,085 individuals, linked 7,122 genetic variants to cell type abundance QTLs (i.e., ccQTLs) of predefined blood cell types, measured using flow cytometry^2,3^. Importantly, these ccQTLs often colocalize with disease^2–4^, providing a mechanism how cell type composition changes may be involved in disease pathogenesis. Moreover, the largest molecular QTL analyses to-date have been conducted using gene expression data, i.e., expression QTL (eQTL) mapping. Such efforts span broad multi-tissue initiatives, as done in the Genotype-Tissue Expression (GTEx) consortium that mapped eQTLs in 838 donors across 49 tissues^5^, as well as large single tissue studies, exemplified by the eQTLGen consortium that mapped eQTLs in whole blood of 43,301 individuals^6^. Despite these large-scale efforts, no potential underlying molecular mechanism has been identified for most disease-associated variants. For example, the GTEx consortium found that only 43% of disease-associated loci colocalize with a known local (*cis*) eQTL effect in at least one tissue^2^. Similarly, the eQTLGen consortium (phase 2) found a *cis* or distal (*trans*) eQTL effect for 45.9% of the 1,848 independent immune disease-associated SNPs tested^6^. Different explanations have been provided for this relatively small overlap. First, many genetic variants may act only in a specific cellular or environmental context^7,8^ that is not adequately captured by tissue-level studies such as GTEx^5^ or eQTLGen^6^. Second, molecular readouts other than gene expression levels (e.g., splicing, chromatin accessibility) may be needed to capture additional overlap with disease-associated variants^9^. Finally, despite the enormous sample sizes of current eQTL consortia, statistical power to detect genome-wide eQTL effects is still limited for tissues other than blood. Yet it is these context-dependent and distal eQTL effects that may be most relevant for our disease understanding^10,11^.

Single-cell RNA-sequencing (scRNA-seq) data provide a unique opportunity to unbiasedly detect high resolution ccQTLs and context-dependent eQTL effects. However, currently available individual single-cell datasets are still limited in sample size compared to previous bulk QTL consortia efforts^8,12,13^. To increase the power of sc-QTL studies, we initiated the sc-eQTLGen consortium in which the scRNA-seq and genotype data from individual cohorts are processed with standardized computational pipelines so that their ccQTL or eQTL summary statistics can ultimately be combined in a federated QTL meta-analysis^14^. In the current study, we unify 12 single-cell peripheral blood mononuclear cell (PBMC) datasets in a federated QTL meta-analysis and map cell type abundance and cell type-specific regulatory variants across 2,032 donors. We then fine-map these effects, integrate them with a large collection of GWAS studies and with *cis-* and *trans-*eQTLs from the eQTLGen consortium, and demonstrate their value for interpreting immune disease risk (**Figure 1**). This information helps pinpoint how and in which cell type these GWAS risk variants act, thereby providing insights that may inform risk stratification, therapeutic development, and prevention strategies.

**Figure 1.**
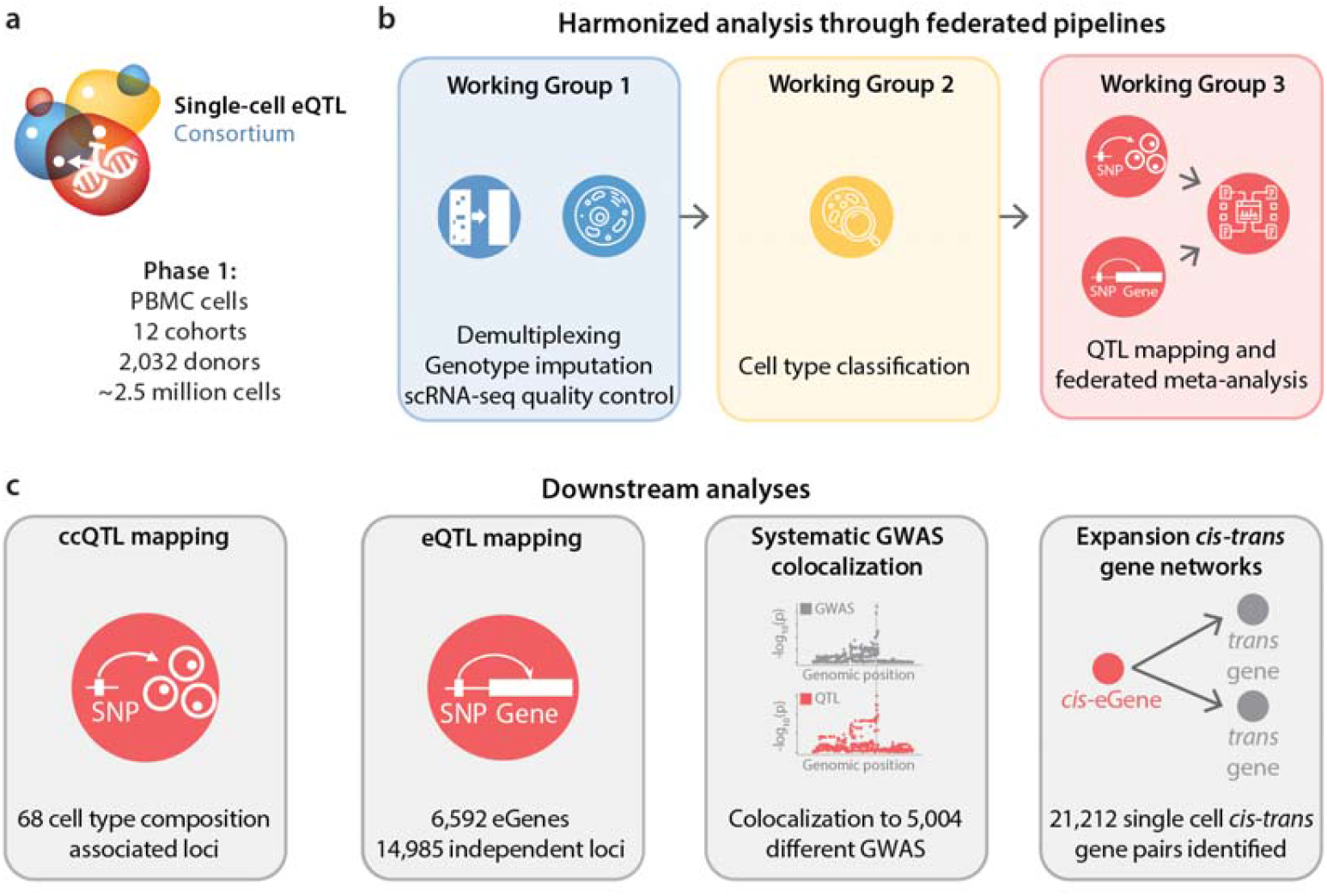
Overview of the single-cell eQTLGen consortium. **a.** We initiated the sc-eQTLGen consortium (Phase 1), which currently includes meta-analysis QTL summary statistics derived from 12 cohorts for 2,032 donors and each of the cell types found in PBMCs. **b**. Standardized computational pipelines were developed to harmonize (WG1) quality control, demultiplexing, (WG2) cell type annotation, and (WG3) QTL mapping across cohorts, enabling centralized meta-analysis of QTL summary statistics. **c**. We conducted several downstream analyses: 1. ccQTL mapping on 63 different cell type composition estimates (on both major PBMC cell types and their subtypes), identifying 68 unique genetic signals associated with differences in cell type proportion; 2. cell type-specific eQTL analysis, which identified 6,592 eGenes and 14,985 independent eQTL effects; 3. systematic colocalization of eQTLs with over 7,000 different GWASs identified colocalization to 5,004 GWASs; 4. systematic colocalization of *trans*-eQTL signals from the bulk blood eQTLGen study (n = 43,301 samples) with our cell type-specific *cis-*eQTL signals, which identified 21,212 *cis-trans* gene pairs.

## Results

### Building harmonized pipelines to perform QTL meta-analysis across 12 cohorts

We performed ccQTL and eQTL meta-analyses in the sc-eQTLGen consortium, combining scRNA-seq data from 2,032 individuals and ∼2.5M PBMCs (**Figure 1a**, and **Table S1**). We harmonized data from 12 cohorts (**Note S2**), generated by different single-cell technologies and chemistries (e.g., full-length Smart-seq2, 3’- and 5’-end 10X Genomics) and from individuals from multiple ancestries, different ages, and both sexes (**Table S1**). Specifically, we included the OneK1K study^13^ (1,017, European ancestry), CD4+ T cells from the Trynka study^15^ (117, European ancestry), CD4+ T cells from the ARMS study^16^ (45, European ancestry), the 300BCG study^17^ (38, European ancestry), the OASIS study^18,19^ (91, East Asian Ancestry), cerebrospinal fluid (CSF) data^20^ from the Sawcer lab (73, European ancestry), the Burkina Faso malaria study^21^ (178, African Ancestry), the Ye SLE study^12^ (170, European, East Asian and Southern Asian ancestries), the van der Wijst study^22^ (40, European ancestry), unpublished single nucleus dataset (118, European ancestry), and unstimulated samples from the 1M-scBloodNL project^8^ (120, European ancestry) and the van Blokland STEMI study^23^ (38, European ancestry) (**Note S2**).

Because QTL analyses involve privacy-sensitive genotype information, raw data could not be centralized. Instead, standardized analysis pipelines were run within each cohort, and only summary statistics for the federated meta-analyses were shared. To facilitate this setup, we established several working groups (WG) to harmonize: WG1. genotype imputation and scRNA-seq data preprocessing, quality control (QC), and donor demultiplexing, WG2. cell type assignment, and WG3. ccQTL and eQTL mapping (**Figure 1b**, see **Methods**). These pipelines are publicly available on Github (**Code Availability**), and several parts of these pipeline elements have been described in detail in separate papers (sample demultiplexing^24^, cell type assignment^25–27^, and eQTL mapping^28^).

In short, we removed cells that were assigned ambiguously or as doublets using a combination of tools that together capture both heterogenic doublets (from different donors) and homogenic doublets (from the same donor)^24^. Additionally, we removed cells that expressed fewer unique molecular identifiers (UMIs) or RNA molecules than the median minus twice the median absolute deviation (MAD) of the entire dataset or whose mitochondrial gene content percentage exceeded the median plus three times the MAD. Next, we annotated the QC-passing cells to seven major cell types, which were further resolved into 27 cell subtypes: B cells (B intermediate, B memory, B naïve, plasmablast), CD4+ T cells (CD4 cytotoxic T, CD4 naïve, CD4 proliferating, CD4 central memory (TCM), CD4 effector memory (TEM), regulatory T cell), CD8+ T cells (CD8 naïve, CD8 proliferating, CD8 TCM, CD8 TEM), monocytes (classical CD14+ monocyte, non-classical CD16+ monocyte), natural killer cells (NKdim, NKbright, NK proliferating) and dendritic cells (AXL+ DC, conventional DC1 (cDC1), cDC2, plasmacytoid DC (pDC)), ILCs (innate lymphoid cells) and three other T cell subtypes (MAIT, dnT, gdT) (**Table S2**). Cell type annotation was performed using hierarchical scPred^25,27^ and Azimuth^26^. We only retained cells that were concordantly classified by both tools at the major cell type resolution. The annotation reference for these tools was trained on joint scRNA-seq and sc-surface protein data (CITE-seq) of human PBMCs as defined previously^26^. These QCed and annotated cells were then used for downstream QTL analyses (**Figure 1c**).

### Single cell−derived ccQTLs capture bulk-derived *trans-*eQTLs

Flow cytometry-based analyses have previously shown the value of ccQTLs to interpret disease-associated variants^2,3^. To achieve higher cellular resolution in characterizing these ccQTLs, we therefore explored the usage of scRNA-seq data. We assessed the effect of genetics on cell type composition by conducting a ccQTL meta-analysis at different cellular resolutions. Per donor we calculated the ratio of cells belonging to: the lymphoid (B, CD4+ T, CD8+ T, NK, ILC) versus myeloid (monocyte, DC) lineage, each of the seven major cell types versus total PBMCs, each of the 27 cell subtypes versus total PBMCs, and cell subtypes versus the major cell type they belong to, totaling 63 phenotypes. As some of our ccQTLs represent cell type fractions within total PBMCs and ccQTLs typically have effect sizes smaller than eQTLs^29^, we limited our meta-analysis to those datasets that were most similar (7 out of 12 datasets). Specifically, we excluded datasets that provided only partial PBMC representation rather than the full PBMC spectrum (the cerebrospinal fluid dataset (Sawcer CSF^20^) and two datasets that contained only a single major cell type (Trynka^15^ and ARMS^16^)), one dataset with fewer than 40 donors (300BCG^17^), and one dataset from multiple ancestries (Ye SLE^12^). Furthermore, we limited our analysis to donors who had at least 350 total PBMCs, resulting in the inclusion of 1,710 donors. We defined Gaussian-normalized cell type ratios as phenotypes and applied a linear mixed model to test associations with genetic variants, testing variants with MAF >5% and Hardy-Weinberg Equilibrium (HWE) P-value > 1×10^−4^ (see **Methods**). The resulting summary statistics were subsequently combined using an inverse-variance weighted meta-analysis, keeping only effects tested in all datasets. To be conservative, given the remaining differences between datasets, we removed variants with a high association heterogeneity (I^2^ > 40%).

We noted an interdependency across many of the tested cell type proportions. For instance, the proportions of lymphocytes to PBMCs were negatively correlated with those for myeloid cells to PBMCs. ccQTL analysis on these proportions would therefore likely yield the same genetic association signals, albeit with opposite effect directions. Therefore, to more accurately count the number of associated genetic loci, we first defined association loci using “Locus breaker”^30^ (see **Methods and Fig SX**). In these loci we performed fine-mapping on each of the ccQTL summary statistics sets, using SuSiE^31^ with the multiple causal variant assumption (see **Methods**). These fine-mapped loci we termed independent loci. We subsequently performed colocalization of the independent loci with coloc^32^ to identify sharedness of genetic signal among cell composition traits. If fine-mapping was not successful, we deemed the loci to be the same if the association regions overlapped, and counted non-overlapping regions as a single independent locus.

After applying fine-mapping and colocalization, we identified three genome-wide significant independent loci (P < 5.0×10^−8^), associated with cell subtype composition in monocytes, CD8+ T cells, and NK cells (**Figure 2a** and **Table S3**). At a suggestive threshold (P < 5.0×10^−6^), we identified an additional 65 independent loci, including loci influencing the proportion of B cells, CD4+ T cells, and DCs (**Table S3**). Thus, in all, we found 68 independent genetic signals associated with cell type composition derived from PBMC single-cell data. Most of these signals corresponded to known regulators of immune cell abundance: 63 of the 68 independent loci (92.6%) overlapped with cell count traits in the UK Biobank (UKB) cell composition GWAS^3^ (leukocyte count, monocyte count, lymphocyte percentage of leukocytes, and monocyte percentage of leukocytes) (Table S3). For example, the UKB ‘monocyte count’ locus rs67122419 (P = 3.2×10^−151^) aligned with our genome-wide signal of CD14+ versus CD16+ monocyte proportions. Likewise, a genome-wide association for CD8+ TEM in total PBMC counts colocalized with the UKB ‘leukocyte count’ locus rs2523579 (P = 4.8×10^−191^). Moreover, a suggestive CD4+ T cell in total PBMC counts signal corresponded to the UKB ‘lymphocyte percentage of leukocytes’ rs4648050 (P = 6.2×10^−44^). However, we note that, while nearly all of the independent loci have been reported before^2,3^, the traits do not precisely match, as the UKB measurements were derived from flow cytometry−based technologies. Therefore, some differences between the association signals may exist due to differences in cell count resolution and the use of different measures to quantify cell count traits. By leveraging our single-cell data, we have added resolution to the existing associations; however, as resolutions differ, we cannot directly replicate effect directions.

**Figure 2.**
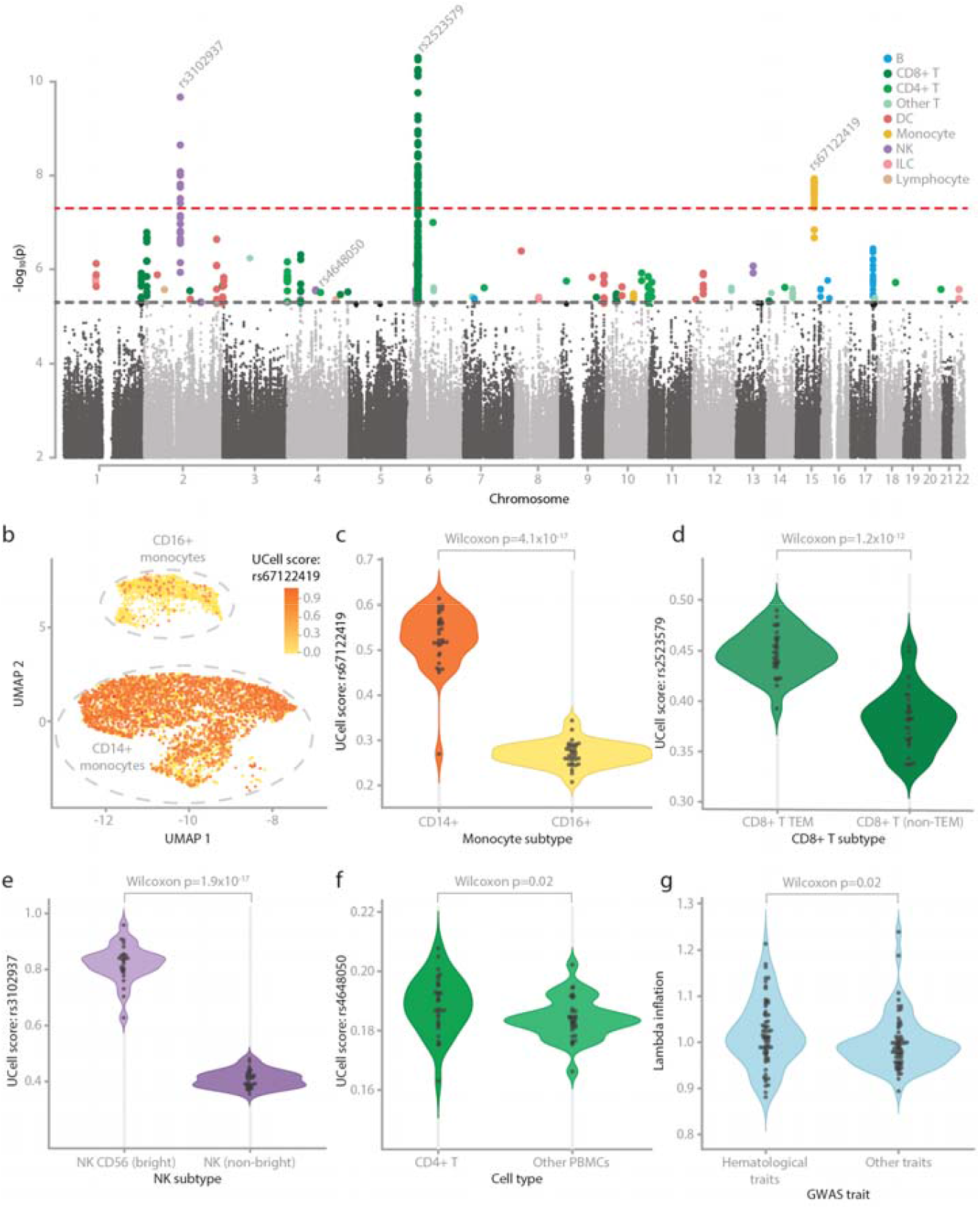
Overview of the ccQTL results. **a.** Combined Manhattan plot of the three genome-wide significant (P < 5×10^−8^) and 65 suggestive (P < 5×10^−6^) cell count GWAS traits. These association signals are colored by the cell type whose proportions are affected by the variant. **b**. Subset of the UMAP for the monocytes. The two cell subtypes are circled (CD16+ on top, CD14+ monocytes bottom). Colors indicate the UCell scores per cell based on the *trans*-eGenes linked to the ccQTL SNP. **c-f**. Violin plots of donor-aggregated averages of the UCell score in the monocyte (**c**), NK (**d**), CD8+ T (**e**) and CD4+ T (**f**) cell (sub)types. **g**. Violin plots of the λ-inflation of the ccQTL variants versus GWAS-associated variants for hematological traits (left) or other GWAS (right) (**Methods**).

As an alternative way to validate the ccQTLs at the appropriate resolution, we reasoned we could use the *trans*-eQTLs identified in bulk data. These *trans-*eQTLs are likely a mixture of genetic effects on cell type composition and intracellular expression regulation. For instance, if a gene is predominantly expressed in a specific cell subtype, this can lead to an apparent *trans*-eQTL effect in bulk RNA-seq data if the associated variant changes the proportion of that cell subtype, rather than changing gene expression. Therefore, we colocalized our ccQTLs with *trans*-eQTLs identified using bulk RNA-seq data from 43,301 whole blood samples in eQTLGen (phase 2)^6^. Then, using one of the representative scRNA-seq datasets (UMCG v3^8,23^) in sc-eQTLGen, we used UCell^33^ to compute a rank-based expression score indicating how highly expressed the ccQTL-associated trans-eGenes are in a single cell (**Methods**). Scores were averaged per cell (sub)type per donor. We next tested whether UCell scores differed between the relevant cell type and the other cells contributing to the fraction underlying the ccQTL phenotype, using a paired Wilcoxon test. This approach allowed us to assess whether the bulk-derived *trans*-eGenes that were associated with the same genetic variant as the ccQTL were differentially expressed in the cell type implicated in the ccQTL. At the same time, it enabled us to distinguish *trans*-eQTL signals driven by changes in cell type composition from those reflecting intracellular regulatory effects.

We could apply this approach to four independent ccQTL loci, as we required the ccQTL to colocalize with at least two *trans-*eGenes: two out of three genome-wide loci (monocyte and NK effects) and two suggestive loci (CD4+ T in PBMC and CD8+ TEM in CD8+ T cells). All four ccQTL loci were validated using this approach: the module scores were significantly different between the cell (sub)types and were highest in the cell type whose abundance increased in association with the specific allele (monocyte locus: Wilcox P = 4.1×10^−17^; NK locus: Wilcox P = 1.9×10^−17^; CD8+ TEM locus: Wilcox P = 1.2×10^−12^; CD4+ T locus: Wilcox P = 0.02) (**Figure 2b−f**). While we expect that our list of ccQTLs and *trans*-eQTLs are incomplete, the concordant replication and consistent effect directions provide additional support for the robustness of our ccQTL findings.

Next, we assessed whether the identified ccQTL loci contribute to disease, as was previously reported for flow cytometry-based ccQTLs^34^. To this end, we examined the lambda (λ) inflation to evaluate whether our ccQTLs show an enriched association signal in external GWASs. We confirmed that our ccQTL variants showed a larger λ-inflation for hematological measurement traits as compared to other GWAS (Wilcox P = 0.012, median λ-inflation difference = 0.02, **Figure 2g**), indicating enrichment for genetic determinants of blood cell composition. In contrast, we observed no λ-inflation for disease or immune system disease in general, or any specific immune disease GWAS—likely reflecting the limited number of significant ccQTL and disease loci. The absence of immune disease-related enrichment compared to previous flow cytometry-based measurements^34^ may reflect the substantially smaller sample size of our study (up to 224-fold fewer individuals) or that disease-relevant effects arise in cell types not represented in PBMCs, such as neutrophils or eosinophils.

### Single-cell data detects novel eQTLs that are more often lineage-specific than bulk-derived eQTLs

Many ccQTLs likely act through *cis*-regulatory mechanisms: a genetic variant first influences the expression level of a specific gene (*cis*-eQTL) involved in, for example, lineage differentiation, after which cell type composition is affected. However, genetic variants can also affect gene expression levels without altering cell type composition. Therefore, to systematically capture such regulatory effects, we conducted a *cis*-eQTL meta-analysis across all 12 cohorts, assessing all QC-passing variants within a 1 megabase (Mb) distance around the gene (see **Methods**). We limited the eQTL analysis to the six most abundant major cell types (CD4+ T, CD8+ T, NK, B, Monocyte and DC), to have sufficient cell numbers in each dataset for reliable pseudobulks. Across all cell types, we detected eQTL effects for 6,592 genes (eGenes), covering 56.4% of the expressed genes (see **Methods**) (**Figure 3a**). Compared to the largest sc-eQTL study within our meta-analysis (OneK1K, n = 1,018 donors), doubling the number of donors through meta-analysis increased the number of detected eGenes by 1.3x (for CD4+ T cells) to 6.7x (for DCs) (**Figure 3a**). The number of detected eGenes per cell type was closely related to the average number of donors and cells per donor available for that cell type (**Figure 3b, Figure SX**): we identified the highest number of eGenes in the cell type for which most donors and cells per donor were available (CD4+ T: 4,988 eGenes, 39.2% of all expressed genes), and the fewest for the cell type with least donors and fewest cells per donor (DCs: 375 eGenes, 3.8% of all expressed genes).

**Figure 3.**
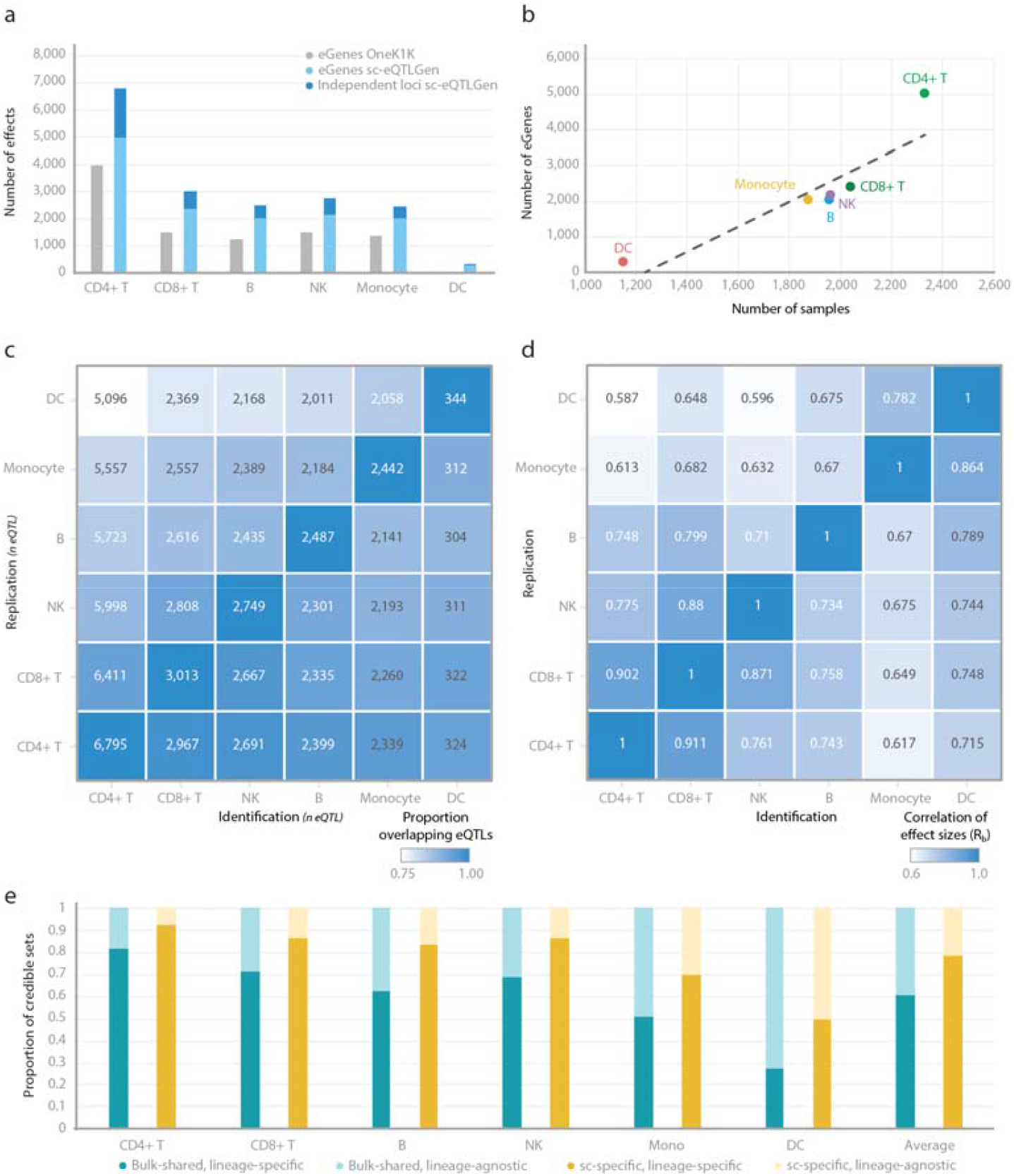
eQTL meta-analysis results. **a.** Comparison of the number of eGenes in sc-eQTLGen (light blue) and the number of eQTLs in the largest individual dataset (OneK1K^13^, gray). Additionally showing the number of independent loci after fine-mapping (darker shade of blue). **b**. Scatter plot showing, per cell type, the average number of cells per donor versus the number of detected eGenes. **c**. Heatmap showing the number of shared expressed genes between the six cell types. Columns are the discovery cell type. Rows are the replication cell types. d. Heatmap as in **(c)** but now showing the r_b_ between the eQTL effect sizes of the shared expressed genes. **e**. Bar chart depicting the number of independent loci observed in the six cell types annotated by their replicability in bulk eQTLGen^6^ (blue = bulk-shared, yellow= sc-specific). The bulk-shared and sc-specific loci are further subdivided by their replicability across blood lineages (darker shade = lineage-specific, lighter = lineage-agnostic).

Multiple genetic variants within a cell type or across cell types may regulate the same eGene. First, to identify the independent loci that regulate the same eGene within a cell type, we conducted fine-mapping using SuSiE^31^. In total, we identified 14,985 independent eQTLs (**Table S4**), increasing the total number of eQTL signals by 20.41⍰ 36.2% (**Figure 3a**). We observed the highest number of independent effects in CD4+ T cells, in line with the higher numbers of donors and cells for that cell type. Most independent eQTL effects were identified within 100 kb around the transcription start site of the gene they are regulating. In line with previous observations^35^, secondary eQTL effects were, on average, further from the gene than the primary eQTL effect (**Figure S1**).

We then studied the cell type-specificity of eQTL effects by comparing the effect of significant eQTLs across cell types using two complementary strategies. In the first, we assessed pair-wise similarity across cell types between expressed genes (**Figure 3c**) and between eQTL effect sizes (**Figure 3d**) using the r metric^36^. In the second strategy, we employed systematic colocalization of the fine-mapped loci across the 6 major cell types and over genes (**Figure 3e**). While the r_b_ metric evaluates whether shared genes are similarly regulated by genetics, the colocalization formally tests whether the genetic signal is truly overlapping, but only for signals that are strong enough to be fine-mappable. The first strategy showed that whether a gene is expressed (**Figure 3c**) and the gene’s genetic regulation (**Figure 3d**) are more similar within cells from the same lineage than across lineages. For instance, CD4+ and CD8+ T cells showed high concordance of eQTL effect sizes (r_b_ = 0.9), whereas the CD4+ T cells and monocytes showed substantially lower concordance (r_b_ = 0.6). The colocalization strategy supported these observations. We identified 6,434 independent loci across the six major cell types, allowing for loci to affect multiple genes (**Figure S2**). Of these, 82.0% were associated with a single gene, although some, particularly in the HLA locus, affected up to 66 genes. Overall, most loci (62.2%) were specific to a single cell type. This is likely in part explained by the relatively large number of *cis*-eGenes uniquely detected in CD4+ T cells due to higher statistical power in this cell type. However, when restricting our analysis to the 431 genes tested in at least five of the six cell types, the results were more similar to those of the r_b_-based approach (**Figure S3**: 91% of these genes were shared within rather than across lineages (85%).

Next, we assessed how many of our independent eQTL loci were also found in eQTLGen (phase 2)^6^ (**Methods**). We observed comparable results for each of the six cell types (**Fig. 3e**). Overall, 57.8% of the independent loci were found in both the sc- and bulk-derived datasets and shared at least one eGene, providing confidence in our results while also allowing annotation of the cellular context in which these genetic variants act. The remaining loci either did not have any overlapping eGene assigned (17.5%) or were not identified in eQTLGen (24.7%). As expected, loci shared across single-cell and bulk data were more often shared across lineages (on average 39.6% cross-lineage effects for bulk-shared effects and 21.0% for single-cell-specific effects, **Figure 3e**). Together, these results highlight the added value of single-cell data for detecting novel, often lineage-specific, eQTLs and for providing the context in which an eQTL acts.

### Single-cell QTLs shed new light on disease

Next, we determined whether and how single-cell QTLs provided new disease insights. As we assumed that all ccQTLs relevant to disease will act through eQTLs, we related all our comparisons to eQTLs first. For this, we colocalized the eQTL loci to GWAS loci by leveraging the OpenTargets Genetics database^37^ of more than 7,000 GWASs. We found 2,271 independent loci (6,434 eQTL, **Figure 4a**) were linked to 5,004 different GWASs (Table S5). Among those GWASs, immune system disease loci were most often colocalizing with eQTL loci (median ratio 0.40) and least often with hematological measurement loci (median ratio 0.30) (**Figure 4b**). Interestingly, other diseases than immune system diseases also showed higher colocalization ratios than hematological measurements (median ratio = 0.34), likely due to some immune involvement in those diseases as well (**Figure 4c**).

**Figure 4.**
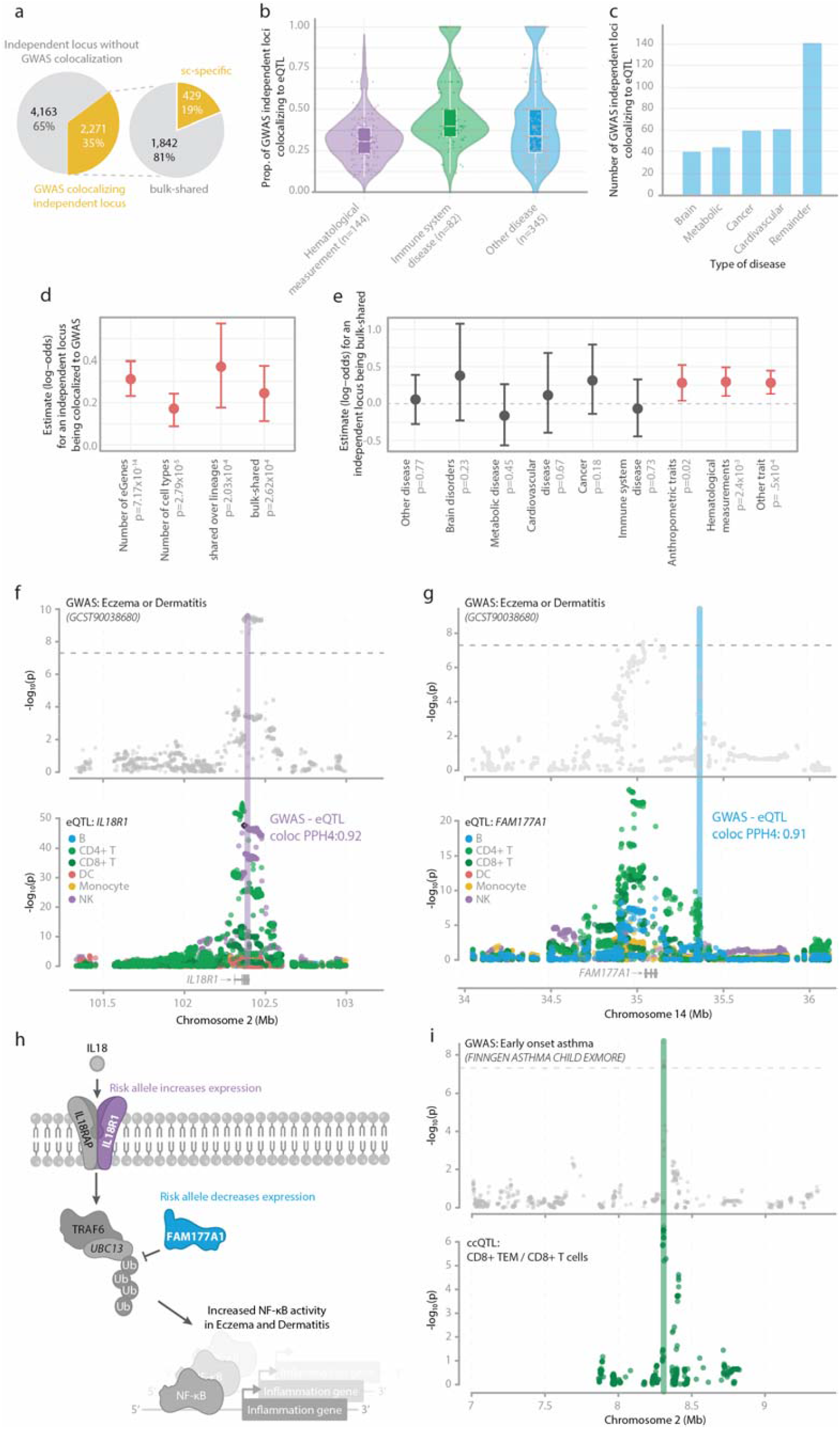
QTL to GWAS trait colocalization. **a**. Systematic colocalization of independent eQTL effects with GWAS traits. For colocalizing signals, sharing of eQTLs between single-cell and bulk eQTLGen was evaluated. **b**. Per GWAS trait category the proportion of genetic signals that colocalized with eQTLs. **c**. The other disease category (from b) split into more precise disease categories, demonstrating that immunelZlderived eQTLs inform traits beyond immune diseases. **d**. SNP features that increase chances (log OR) that a sc-eQTL colocalizes with GWAS traits. e. Likelihood (log OR) that eQTL loci shared between single-cell and bulk eQTLGen, colocalize with different GWAS traits (black: not significant, red: significant). **f**. & **g**. Example locus plots for loci that are uniquely associated to sc-eQTL effects colocalizing to the GWAS on “Eczema or dermatitis”. **f**. Locus plot for the colocalization between the GWAS signal (rs12712145) and *IL18R1* in NK cells. **g**. Locus plot for the colocalization between the GWAS signal (rs28618323) and *FAM177A1* in B cells. **h**. Illustration of the hypothetical mechanism by which IL-18 signaling to NF-kB plays a role in the disease pathogenesis of Eczema and Dermatitis. **i**. Example of disease association (early onset of asthma) identified by ccQTL effect in sc-eQTLGen.

To better understand which eQTL loci are more likely to colocalize to GWAS loci, we used logistic regression (see **Methods**). This revealed that eQTL loci affecting a larger number of genes (P = 7.2×10^−14^), detected across multiple cell types (P= 2.8×10^−5^), or identified in both lymphoid and myeloid cells (P= 2.0×10^−4^), were significantly more likely to colocalize with a GWAS signal (Figure 4d). Interestingly, these same analyses showed that eQTL loci shared between single-cell and bulk eQTLGen are more likely to colocalize to GWAS traits (P= 2.6×10^−4^) (**Figure 4d**). More specifically, when assessing characteristics of shared loci we found that these are significantly enriched in hematological measurements (P= 2.4×10^−3^), anthropometric traits (P= 0.02), other (non-disease-associated) traits (P= 5.0×10^−3^) (**Figure 4e**). These colocalization-based observations are in line with SNP-based enrichments (considering linkage disequilibrium (LD)), in which both the immune system disease GWAS (EFO0000540, cross cell type Fisher OR: 8.7, P= 1.6×10^−48^) and all disease trait GWAS sets (EFO0000408, cross cell type Fisher OR 4.1, P= 8.1×10^−104^) were enriched in our sc-eQTLs over all other SNPs assessed (**Table S6**). Notably, we observed that the sc-eQTLs are significantly more enriched for disease GWAS than bulk eQTLGen eQTLs (Z-test OR 2.9, P= 4×10^−4^, **Table S6**), underscoring the added value of sc-eQTLs for disease interpretation. Together, these analyses show that effects unique to single-cell data are more relevant to the interpretation of disease GWAS loci, and shared effects are more likely to capture cell (sub)type composition related loci.

As an example of a disease association uniquely identified using sc-eQTLs, we considered two eQTL loci that both colocalized with ‘eczema or dermatitis’ (GCST90038680)^38^. The first locus, containing risk variant rs12712145 (Chr2:102,392,150, GWAS P = 2.2×10^−10^), was present in NK and CD4+ T cells, but different genes were implicated depending on the cell type. The C risk allele increased expression of *IL18R1* in NK cells (eQTL Z = 14.6) but lowered expression of *IL18RAP* in CD4+ T cells (eQTL Z = −0.45, **Figure S4**). The second locus was found in B cells, where the disease risk allele A of variant rs28618323 (Chr14:35,106,586, GWAS P = 8.2×10^−10^) decreased expression of *FAM177A1* (eQTL Z = −6.1). Together, these genes are all involved in the same signaling pathway. IL18R1 and IL18RAP form the IL18 receptor complex that activates downstream NF-κB signaling upon binding of IL18^39^. FAM177A1 competitively binds TRAF6, preventing its downstream activation of NF-κB signaling^40^. This implies that the elevated IL-18 signaling in atopic dermatitis patients, which also correlates with disease severity^41^, may be facilitated through increased signaling in NK cells and reduced suppression in B cells (**Figure 4h**).

Another locus uniquely identified in single-cell data as both ccQTL and eQTL and associated with disease (early onset of asthma, FINNGEN_R12_ASTHMA_CHILD_ EXMORE, GWAS P = 5.5×10^−9^), was the risk variant rs391934 (Chr2:8,304,364, **Figure 4i**). The T risk allele increased the proportion of CD8+ TEM to total CD8+ T cells (ccQTL Z-score = 4.8) and increased the expression of the long-non-coding RNA *LINC00299* in NK cells (eQTL Z-score = 4.9, **Figure S5**). Although CD8+ TEM have not been directly associated with asthma before, CD8+ TEMRA have been^42^. This terminally-differentiated subset of CD8+ T cells can arise from CD8+ TEM that re-express CD45RA^43^. Because our eQTL meta-analyses were conducted primarily on 3’-end scRNA-seq data, we could not readily distinguish the CD8+ TEM and CD8+ TEMRA subtypes, which are defined by their expression of different CD45 protein isoforms (CD45RO/CD45RA) encoded by the *PTPRC* gene. We therefore suspect that this ccQTL may have instead captured a change in the abundance of CD8+ TEMRA cells. Further research is needed to define whether the regulation of *LINC00299* expression in NK cells contributes to the abundance change of the CD8+ TEMRA subset, or that the eQTL effect may have been present in the CD8+ TEM subset as well as a consequence of the ccQTL. Notably, the latter possibility is supported by the high expression of *LINC00299* within the CD8+ TEM subset.

### Expanding disease-relevant cis-trans gene pairs identified in bulk by leveraging single-cell resolution

Finally, we assessed whether the *cis*-eQTLs unique to single-cell data could also help refine gene regulatory networks relevant to disease. Ideally, this would involve mapping these *cis*-eQTLs and *trans*-eQTLs in our single-cell data, such that *cis*-regulatory variants can be directly linked to their downstream *trans* effects. However, the sample size of our single-cell meta-analysis is underpowered for genome-wide *trans*-eQTL discovery. In contrast, eQTLGen (phase 2)^6^ provides adequate power for such analyses, with a sample size >20-fold larger. Within eQTLGen, 48,037 *trans*-eQTLs were identified (defined as effects >5Mb away from the eSNP, reflecting 9,199 unique trans-eGenes), with 23,248 (52.8%) linked to at least one *cis*-eGene (**Table S7**). These *cis*-*trans* associations enabled inference of directional gene-gene relationships, offering additional insights into disease mechanisms. To leverage both the resolution of the single-cell data and the statistical power of bulk data, we integrated these approaches to improve identification of true regulatory *cis*-*trans* gene pairs associated with disease.

First, we examined how many sc-derived *cis*-eQTLs could be colocalized with bulk-derived *trans*-eQTLs: 14,683 *trans*-eQTLs (5,563 *trans*-eGenes). Most variants could not be colocalized as there was no *cis*-eQTL effect in the single-cell data. In total, we identified 6,382 *trans-*eQTLs that colocalized with at least one sc-derived *cis*-eQTL, corresponding to 3,706 *trans*-eGenes. 37.2% of these colocalizing *trans*-eQTLs (**Figure 5a**) were newly annotated to at least one *cis*-eQTL using the single-cell data and most of these links (58%) were associated with disease (annotation based on eQTLGen^6^). In total these colocalizing effects reflected 21,212 unique *cis-trans* gene pairs (linking on average 5.7 *cis*-eGenes to a *trans*-eGene), including 15,563 (73.4%) that were not observed in eQTLGen.

**Figure 5.**
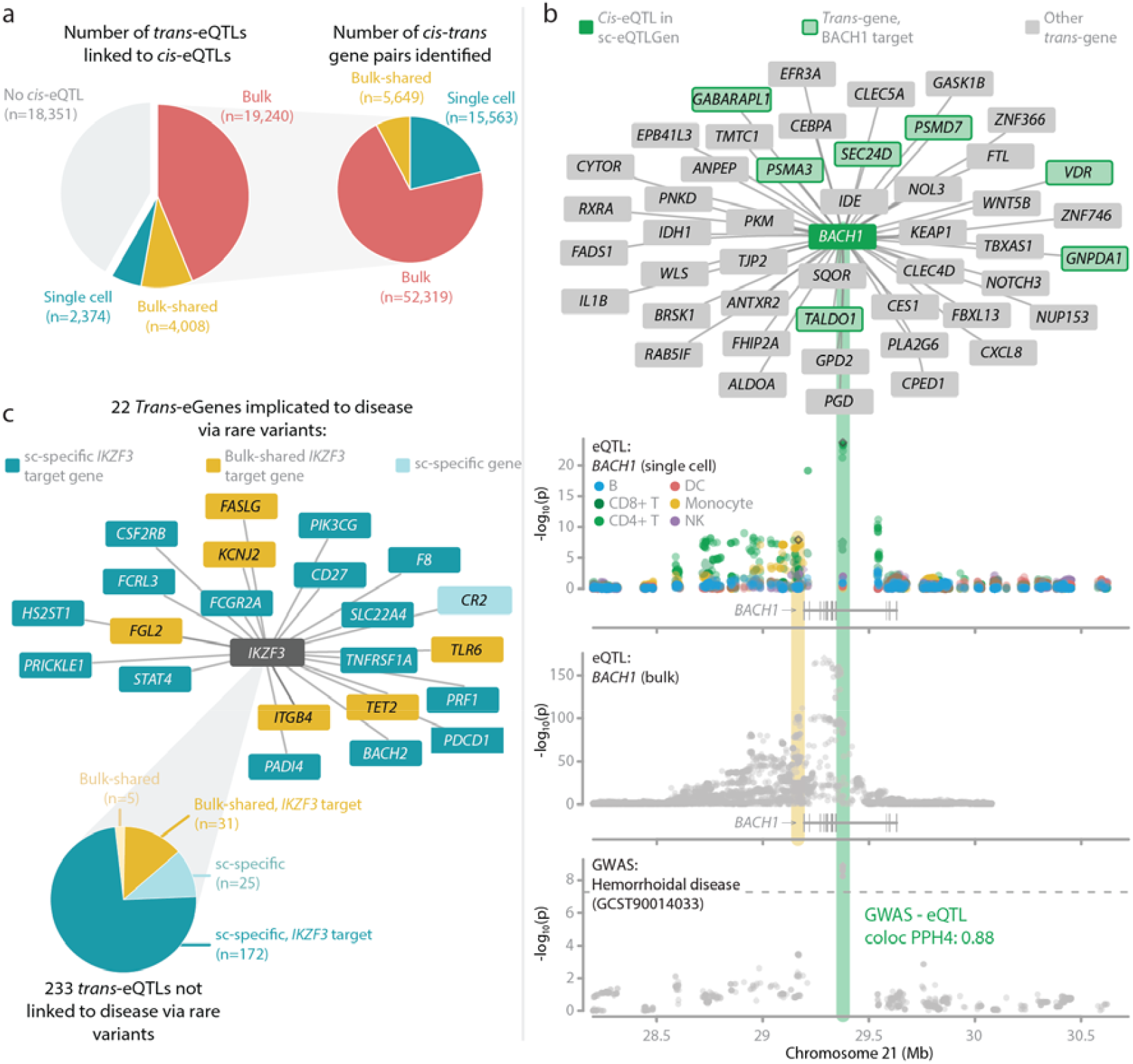
Distal disease networks uncovered via single-cell eQTL mapping. **a**. The number of bulk eQTLGen^3^ *trans*-eQTLs that colocalize to single-cell or bulk eQTLGen *cis-*eQTLs, and the number of directional *cis-trans* gene pairs that could be inferred from that. b. Single-cell-specific *cis-trans* gene pairs identified for the T-cell-specific, but not monocyte *cis*-eQTL locus for *BACH1*. Only the T-cell-specific *BACH1 cis*-eQTL locus colocalizes with hemorrhoidal disease, providing mechanistic insights into how this locus contributes to the disease pathogenesis. Importantly, these effects could not be identified in bulk as the bulk-averaged *cis*-eQTL signal did not colocalize to either the *trans-*eQTLs or the disease GWAS. c. *Cis-trans* gene pairs identified for the bulk and single-cell *IKZF3 cis*-eQTL locus. Many additional *trans-*eGenes were linked to the *IKZF3 cis*-eGene using single-cell eQTL data. These additional *trans*-eGenes were enriched for *IKZF3* regulation, and a subset was also involved in rare disease through previously reported rare variants.

A particularly noteworthy single-cell-specific effect was a *trans*-eQTL locus linked to a *cis*-eQTL locus for *BACH1*. Two independent *cis*-eQTLs were found in single-cell data: one in CD4+ and CD8+ T cells (rs2832300) and the other in monocytes and CD8+ T cells (rs66723060) (**Figure 5b**). Interestingly, only the T cell-specific locus was linked to the *trans*-eQTL effects, corresponding to 45 *trans-*eGenes. These genes were significantly enriched for *BACH1*^43^ P = 0.01 (corrected for the number of assessed sets, **Methods**)), supporting the validity of the observed *trans*-eQTL effects. The *cis*-locus was further implicated in a GWAS for hemorrhoidal disease^44^ (PP H4: 0.88), a condition influenced by genetic, immunological, and metabolic factors^30^. Consistent with this, pathways related to the immune system (Reactome:R-HSA-168256, g:Profiler P = 5.4×10^−5^) and metabolism (Reactome:R-HSA-1430728, g:Profiler P = 2.7×10^−5^) were enriched among the *trans*-eGenes after accounting for genes expressed in eQTLGen. The T cell-specific locus was not detected in the bulk data, likely due to the averaging of the two sc-eQTL loci in bulk data (**Figure 5b**). This was supported by the higher LD between bulk (rs10563577) and single-cell lead variants (R^2^ = 0.29 bulk-monocyte effect (rs66723060); R^2^ = 0.37 bulk-CD4+ T cell effect (rs2832300)) than between the two single-cell effects (R^2^ = 0.12 monocyte-CD4+ T cell).

Moreover, even for the *trans*-eGenes previously annotated using bulk data, single-cell *cis*-eQTLs provide additional disease insights. For example, in bulk eQTLGen data, the *IKZF3 cis*-eQTL colocalized with 48 *trans*-eGenes, whereas leveraging sc-eQTLGen data allowed us to annotate the *IKZF3 cis*-eQTL as lymphocyte-specific and link the SNP to 255 *trans*-eGenes. Using also these additional *trans*-eGenes, we found *IKZF* motifs to be significantly enriched among the linked *trans-* eGenes (g:Profiler P = 3.0×10^−3^, precision = 87%), providing additional support for *IKZF3* being the key regulator of these genes. Interestingly, the *IKZF3 cis*-eQTL locus colocalized with several autoimmune and allergic disease GWASs, including asthma (PP H4: 0.97), Crohn’s disease (PP H4: 0.99), and rheumatoid arthritis (PP H4: 0.97). Additional evidence for the role of the 255 *trans*-eGenes in these diseases was provided by a subset of 22 rare variant-implicated disease genes that were also previously reported in Orphanet, ClinVar, or gene-burden analyses, as collected within the Open Targets genetics platform^37^ (**Figure 5c**). Importantly, 15 of these genes were only linked through single-cell data, further strengthening the connection with *IKZF3* and supporting the role of this locus in regulation of genes critical for disease in general. These findings highlight the value of integrating single-cell and bulk eQTLs to identify putatively causal *cis-*to-*trans* eQTL pairs that can help uncover disease-relevant gene regulatory networks.

## Discussion

We performed a ccQTL and eQTL meta-analysis in scRNA-seq data from up to 2,032 individuals in 2.5M peripheral immune cells. This meta-analysis setup increased the donor and cell sample size by more than two-fold compared to the largest study within the consortium^13^, while also covering a wider range of different ancestries (African, European, East Asian and South Asian). Our ccQTL analyses identified three genome-wide significant and 65 suggestive CSs that affected the cell subtype composition of blood. Although most identified loci had previously been implicated with blood composition effects^2,3^, our single-cell framework resolves these effects at substantial higher cellular resolution, revealing which specific immune populations are genetically perturbed. In addition, we could validate our findings by overlaying our ccQTL results with whole blood bulk-derived *trans*-eQTLs from over 43 thousand individuals^6^. The module scores of the *trans*-eGenes that share genetic signal with our sc-derived ccQTLs differed most in the cell type affected by the ccQTL, indicating correct assignment of a composition effect. This also shows that these specific *trans*-eQTLs, i.e., those linked to ccQTLs, likely reflect cell composition effects rather than intracellular expression effects. These results showcase how combining sc-and bulk-derived QTL results can synergize for interpretation of ccQTL and *trans*-eQTL effects. Moreover, the large number of replicated effects we observe, including the suggestive loci, indicates that many more reliable ccQTLs are expected to be discovered with increasing sample size.

Besides genetic effects on blood cell proportions, our sc-eQTL meta-analysis substantially expands the catalog of variants known to change gene expression in specific immune cell types. Compared to the largest sc-eQTL study included in our meta-analysis^13^, which included about half the donors and one-third of the total number of cells, we discovered between 25-363% more *cis*-eQTLs, showcasing the value of our meta-analysis approach. We observed that 68% of these eQTLs act in a lineage-specific manner, and it was this category that was enriched among the 42% of eQTLs undetected in bulk blood-based eQTL meta-analyses efforts, even in over 20-fold more individuals^6^. This underscores a key limitation of bulk analyses: averaging across heterogeneous cell populations dilutes regulatory effects that are confined to specific immune lineages or activation states. As it is exactly these cell- and context-specific eQTLs that are more strongly enriched for disease^44^, this explains why stronger enrichment for GWAS loci is observed among our single-cell-specific eQTLs as opposed to those shared with bulk data.

A further conceptual advance of this work lies in integrating single-cell *cis*-eQTLs with bulk *trans*-eQTLs to reconstruct disease-relevant regulatory cascades. While *trans*-eQTL discovery remains underpowered in single-cell data, the large-scale eQTLGen resource^6^ provides a complementary opportunity to map downstream regulatory effects. By linking bulk *trans*-eQTLs to upstream regulators (i.e., *cis*-eQTLs) identified at single-cell resolution, we uncovered thousands of *cis–trans* gene pairs that were previously unannotated in bulk data. This integration not only increased the number of inferred regulatory relationships but also resolved their cellular context, as illustrated by the *BACH1* and *IKZF3* loci, where single-cell *cis*-eQTLs revealed lineage-specific regulatory hubs linked to immune and metabolic disease pathways.

Altogether, our study provides a valuable resource for anyone interested in understanding how genetic variation may impact disease, by identifying new *cis*-eGenes linked to disease, connecting these *cis*-eGenes with *trans*-eGenes to identify the regulatory intermediates that tie them together, and uncovering the cell types in which these variants act. However, several limitations should be considered. First, QTL discovery power and the accuracy of causal variant fine-mapping differ among the cell types, as the total number of cells and cells per donor used for QTL mapping differed widely among the 6 major cell types assessed. These differences across cell types will likely impact our ability to correctly assign cell type-specificity of an eQTL. Therefore, to overcome this issue, instead we assessed the sharedness of eQTL effects at lineage rather than cell type level, as we expected that eQTL effects are most likely shared within the same lineage (lymphoid or myeloid)^45^. Second, technical heterogeneity across cohorts (such as differences in PBMC isolation, preservation, and preparation) likely introduced variability in cell type composition that may exceed the effect size of individual genetic variants. While this variability could have reduced power for ccQTL detection, the federated meta-analysis design and stringent heterogeneity filtering make spurious associations unlikely. Third, some ccQTLs may indirectly affect expression, resulting in an eQTL effect entirely driven by changes in cell (sub)type composition^29^. While this is mainly a risk for bulk eQTL studies, which use samples composed of many different cell types^6,46^, our sc-eQTL meta-analyses may have been impacted by similar cell subtype changes. As a potential example, we described the ccQTL on CD8+ TEM over total CD8+ T cells that also resulted in an eQTL effect on *LINC00299*, a long non-coding RNA primarily expressed in the CD8+ TEM subtype. To address this, future efforts of the consortium will further expand the donor and cell sample sizes to facilitate sufficient discovery power to map eQTLs at higher cell subtype resolution. Finally, while the current study mapped eQTLs at cell type resolution, several recent sc-QTL studies have shown that additional eQTLs can be identified when leveraging cell subtypes or dynamic cell state annotations instead^29,47^. This is a unique aspect of single-cell data that could provide additional resolution over previous bulk efforts, and it would greatly benefit from the large amount of single-cell data available within the sc-eQTLGen consortium. However, this is an aspect we would like to address in future consortium efforts, as the federated meta-analysis setup complicates uniform assignment of cells along a dynamic trajectory.

In summary, this work highlights the benefits of integrating many sc-eQTL studies within a unified meta-analysis framework. Such framework not only improves the QTL discovery power, but also eliminates the need to share privacy-sensitive data or to rerun analyses whenever new datasets are included. We expect that this work will benefit from further increases in sample size in the near future and from combining the discovery power of bulk eQTL data with the cellular and context-specific resolution of sc-eQTL data through the combined efforts of the eQTLGen^6^ and sc-eQTLGen consortia^14^. This will help us interpret disease-associated genetic variation and aid in the implementation of personalized disease prevention and treatment.

## Methods

### Datasets

Data from 12 cohorts was meta-analyzed using harmonized computational pipelines (see below). The data derived from these cohorts is generated using diverse single-cell technologies and chemistries— including full-length Smart-seq2, as well as 3⍰- and 5⍰-end 10x Genomics scRNA-seq and sc-multiome platforms—and spanning individuals of multiple ancestries, age groups, and both sexes (**Table S1**). Detailed information on the data generation of the scRNA-seq and genotype information is provided in the **Supplementary Information**. Notably, scRNA-seq alignment to the reference genome was not harmonized across the studies: all the 10X studies followed (different versions of) the 10X Genomics CellRanger pipeline, while details of alignment of the Smart-seq2 data of the ARMS cohort can be found in the original publication^16^.

### Genotype imputation

Genotype imputation was carried out for each dataset separately on GRCh38 using a Snakemake (v6.4.1)^48^ pipeline. Briefly, sample quality and consistency were ensured by removing samples missing a large amount of genotype calls (call rate >0.9) and ensuring consistency between genotype-determined and self-reported sex (when provided). If necessary, genotypes were lifted over to build 38, using CrossMap v0.4.1^49^. Next, donor genomes were identified to be most similar to African, Native American, European, East Asian or South Asian ancestries using 1000 Genomes samples as a reference^49^. Samples were grouped by genetic ancestry and imputed in each group separately, leveraging the 30X WGS 1000G reference panel^49^. Post imputation genotype quality was ensured by removing variants with a call rate < 0.99, and a MACH R2 quality score < 0.6. For QTL mapping we removed multiallelic variants, and per study we removed SNPs that had a MAF < 1%, had a HWE P < 1×10^−4^ and ensured that every genotype was observed in every study at least once. At the meta-analysis stage, all SNPs with a MAF < 5% over the whole study were removed, leveraging information calculated using Genotype-IO (https://github.com/molgenis/systemsgenetics/wiki/Genotype-IO). Sample and genotype filtering were carried out using plink 1.9 (v1.90b6.24) and plink 2 (v2.00a3LM)^50^, bcftools (v1.10.2)^51^, vcftools (v0.1.17)^52^, Genotype Harmonizer (v1.4.23)^53^, and R (v3.6.3). Eagle (v2.4.1)^54^ was used to phase the genotypes before imputation with Minimac4 (v1.0.2)^55^ using the 1000 Genomes 30x WGS dataset as imputation reference.

### Demultiplexing and Doublet Removal

Souporcell (v2)^56^ and Demuxlet^57^ were used to demultiplex pools that contained multiple unrelated donors and identify heterotypic doublets. The transcription-based doublet detecting methods DoubletDetection (v3.0)^58^, scds (v1.1.2)^59^, and scrublet (v0.2.1)^60^ were additionally used to identify homotypic doublets (those from the same donors). Droplets were classified as singlets if all software packages identified the droplet as a singlet and both Demuxlet and Souporcell annotated the droplet to the same donor.

### Cell type assignment

We predicted the cell type label of the cells using two different methods: Azimuth^26^ and HierscPred (v1.9.2)^25,27^. For Azimuth, we followed the multimodal reference mapping vignette from Seurat (v4.0.5) (https://satijalab.org/seurat/articles/multimodal_reference_mapping). HierscPred is the hierarchical version of scPred. It trains a scPred classifier for every parent node in the hierarchy. During the prediction step, HierscPred starts at the root node and uses the corresponding scPred model to predict which child node the cell belongs to. If the probability of the predicted child node is higher than the threshold (default = 0, so no rejection option), we continued with making predictions using the child node’s scPred classifier. We continued until we reach a leaf node or the prediction probability fell below the threshold. In the latter case, the assigned label is the label of the cell type we last predicted with sufficient confidence. Both methods use the Human-PBMC reference from Hao et al. as the reference/training dataset^14^. This dataset is annotated at three resolutions (celltype.L1, celltype.L2, and celltype.L3). Using both methods, we annotated the cells at the celltype.L2 resolution. Then, based on this L2 resolution, we manually assigned cells to their respective L1 resolution. This L1 resolution was used as input for the eQTL analysis in 6 major cell types: B cells (B intermediate, B memory, B naïve, plasmablast), CD4+ T cells (CD4 CTL, CD4 naïve, CD4 proliferating, CD4 TCM, CD4 TEM, Treg), CD8+ T cells (CD8 naïve, CD8 proliferating, CD8 TCM, CD8 TEM), monocytes (CD14 mono, CD16 mono), NK cells (NKdim, NKbright, NK proliferating), and dendritic cells (ASDC, cDC1, cDC2, pDC). Since both methods are subject to methodological limitations, we only used cells for downstream analyses when the predictions were concordant at this L1 resolution. For the ccQTL mapping we took the L2 annotations as derived from Azimuth as the L2 label, and additionally included ILCs (L1), as well as the three other T cell groups (MAIT, gdT, dnT). As most datasets were sequenced in batches, we tried to ensure that batch effects did not affect the results by annotating the cells per batch. A singularity image to annotate the cells using both methods is available here: https://github.com/sc-eQTLgen-consortium/WG2-pipeline-classification/tree/main.

### scRNA-seq data preprocessing and quality control

We removed those cells that expressed fewer UMIs or RNA molecules than the median minus 2 times the MAD of the entire dataset or had mitochondrial gene content percentage was larger than the median plus 3 times the MAD. We also removed cells that were assigned ambiguously or as doublet using a combination of tools, see **Demultiplexing and Doublet Removal**. Finally, we kept those cells that were concordantly annotated with both hierarchical scPred^25,27^ and Azimuth^26^ trained on human PBMC reference data, as defined previously^26^, see **Cell type assignment**. After QC, we leveraged the PFlog1PF normalization^61^ to normalize the scRNA-seq data per cell type.

### Linkage disequilibrium calculations

LD was calculated for each dataset separately using the Python (v3.8.12) implementation of LDSTORE2 v2.0_x86_64^62^. For this, we generated the following files per chromosome: bgen, bgi, z, sample, master, and bdose, as described in the documentation for LDSTORE2. We then calculated LD in windows to reduce the overlap in the LD matrices, with the window size determined by the distance between genes. We used a minimum window size of 5M bases, with the start of the window 1M bases before the start of a gene and the end of the window 1M bases after a gene. If there were other genes located within a window, the end of the window was determined by the gene that is furthest along in the window.

To store and share LD efficiently, we performed a principal component analysis (PCA) on the LD matrix to reduce the dimensions of the matrix from variants x variants to samples-1 x variants. This reduced matrix, as well as the components and means, were saved to allow for reconstruction of the LD matrix at the central analysis site, where we selected the number of components to be n samples minus one as this allowed for a near perfect reconstruction, at a fraction of the space of the original LD matrices.

### Cell type composition QTL mapping

For ccQTL analyses, we included all cells that were concordantly annotated to the L1 level, and did not perform any additional QC on the cells. Moreover, we removed donors based on a PCA on the genotype information, excluding genetic outliers based on 2x SD within the respective study. We assessed the effect of genetics on different compositional resolutions: 1. lymphoid versus myeloid lineage, 2. major cell type compared to total PBMCs, 3. minor cell type compared to total PMBCs, and 4. compared to the major cell type it belongs to. We limited our meta-analysis to 7 out of the 12 datasets, as we excluded the cerebrospinal fluid dataset (Sawcer CSF), one dataset with fewer than 40 donors (300BCG), datasets from multiple ancestries (Ye), and those that contained only a single cell type (Trynka and ARMS). In the remaining datasets, we limited our analyses to donors that had at least 350 PBMCs. We mapped ccQTLs by running a linear mixed effect model, implemented in LIMIX QTL^28^, on cell count ratios forced to a normal distribution (Gaussian-normalized), which we corrected for age, sex, and total number of cells per donor (n_cells) by including these as fixed effects in the linear model, and accounted for possible population substructure and repeated observations from the same donor by including a PLINK^50^ king kinship matrix (u_K_) as a random effect as follows:

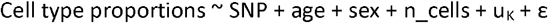

### eQTL mapping

All 12 datasets were used for eQTL mapping. Pseudobulk mean averaged expression data per cell type (L1 resolution) and per donor was used as input for the eQTL mapping^28^. We removed donors whose gene expression (Gaussian-normalized) deviatedmore than two times the SD from the mean of both PC1 and PC2 within the respective study. For the Ye dataset, we deviated from this as this dataset had samples from two ancestries (European and Asian). For this dataset, we included the first two genetic PCs as additional covariates when mapping eQTLs. eQTL mapping was performed using LIMIX QTL on Gaussian-normalized expression data, we included 10 expression PCs as fixed effects and added the PLINK king kinship matrix (u_K_) as a random effect.

LIMIX eQTL model:

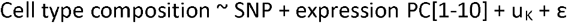

For the Ye dataset:

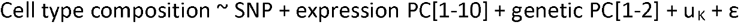

Each dataset performed 100 permutations for the eQTL mapping, after which the permuted and eQTL data were separately meta-analyzed at the central analysis site. To efficiently share the permutation information we calculated Z-scores of the associations (beta/beta_se) for the permuted associations, and fitted a normal distribution over these per SNP. The mean and variance of each association were shared to allow further SNP selection at the central analysis site. At the central site the permutation where combined. Specifically, we sampled from the distributions per association, leveraging the in sample LD, to get back to a full permutation matrix per study. These were meta-anlyzed as outlined in the meta-analysis section. Per gene, per permutation round we selected the top associated variant, and leveraged the beta-distribution to increase resolution when calculating the permuted P value per association as outlined in the FastQTL paper^63^.

### Meta-analysis

Both the ccQTLs and the eQTLs were meta-analyzed at the central analysis site. Here, final SNP and trait selection was performed over all datasets at once (see genotype QC). For ccQTL meta-analysis no further filtering was needed as all traits were shared over the datasets. However, for eQTL meta-analysis, we first subsetted our results to genes expressed in all 12 datasets. For the ccQTL meta-analysis, we used an inverse variance based meta-analysis, the top performing meta-analysis in our benchmarks^64^. In contrast, for the eQTL meta-analysis a sample-size-based weighted meta-analysis^64^ was used instead, as this significantly reduced the amount of data that had to be shared between the sites (inverse variance weighting requires sharing of standard errors per test). To filter associations that are heterogeneous across the datasets we applied the I2 variation filter^65^ after ccQTL mapping, removing effects with I2 over 40 percent.

### Fine-mapping and colocalization analysis

SuSiE was used for fine-mapping^31^, setting the maximum number of credible sets per eGene to 5, leaving the other values at default. For traits without a specific genomic location associated to it, i.e. GWAS traits or ccQTLs, we leveraged the Locus Breaker methodology^30^ to define loci that are used as input for fine-mapping. Throughout the manuscript we refer to these credible sets as independent loci. When SuSiE failed to converge, we defaulted back to reporting a single independent locus per eGene and these effects were not taken forward for colocalizations. We used in sample LD for this procedure, to derive this we meta-analyzed the dataset-specific LD by calculating the sample size-weighted mean LD over the datasets. For the eQTLGen (phase 2)^6^ *cis*-eQTLs we leveraged the same procedure, where we subsetted the eQTLGen summary statistics down to the variants considered in sc-eQTLGen, and used the in sample LD of sc-eQTLGen. This limited differences between the finemapping results for the downstream colocalization. For *trans*-eQTL colocalization, we leveraged the results as taken from eQTLGen to match the disease colocalization in eQTLGen and limit the computational burden due to re-finemapping. To colocalize *cis*-eQTLs to each other, as well as for colocalizing *cis*-*trans* effects we used the coloc package^32^, in which the log bayes factor (LBF) information from SuSiE was used as input, specifically we used the function: “coloc.bf_bf” (from coloc), leveraging default parameters.

To colocalize the sc-eQTLGen *cis*-eQTLs to the Open Targets data we converted the SuSiE^31^ output into StudyLocus format from gentropy v2.0 (https://github.com/opentargets/gentropy) to be consistent with the GWAS credible data format in Open Targets. These independent loci were then overlapped with all GWAS independent loci in the Open Targets ecosystem (March 2025 data release). We considered overlap when two independent loci (GWAS vs eQTL) share at least one variant. The LBFs for each variant were then used to calculate coloc posterior probabilities, where we infer an LBF of 0 for variants which do not exist in one of the two independent loci of an overlap. All analyses were performed on Google Cloud Platform via Airflow using our pipeline orchestration. The mathematics in the coloc implementation is the same as the original coloc R package^32^, and is previously benchmarked against that to ensure it produces the same output. Independent loci were filtered such that the minimal LBF was 2, or >= 0.8686 in log10BF scale, and the independent loci had at least one association at P ≤ 5.0 x 10^−3^. For all colocalization analyses we defined traits to be colocalized if the sum of PP H3 and PP H4 was higher than 0.9 and the ratio between PPH4 and PPH3 was higher than 3.

### ccQTL validation

We validated the ccQTLs by looking up the variants within the ccQTL CSs with UK Biobank cell count GWASes^3^ and whole blood *trans*-eQTLs from eQTLGen (phase 2)^6^. For the latter, we calculated a UCell^33^ module activity score per cell (in the UMCG V3 dataset as representative example) for each independent ccQTL locus. First we colocalized (see **finemapping and colocalization**) each ccQTL locus with *trans*-eQTLs derived from eQTLGen. Next we split up the colocalizing *trans*-eQTLs based on the effect sign, and derived two UCell module scores^33^, one for the positive *trans*-eQTLs and one for the negative *trans*-eQTLs. We subtracted the module score of negatively associated *trans*-eGenes from that of positively associated ones, to get to the final UCell score. For this calculation we required at least two colocalizing trans-eQTL effects per ccQTL loci. We then used a Wilcoxon Signed-Rank test to define whether this module activity score was significantly different in the cells that changed composition relative to all other cells over which the ccQTL was calculated (i.e. versus all PBMCs, or versus the other cells within the same major group depending on the effect), and we assessed whether the direction of effect was the same as the ccQTL effect.

### ccQTL inflation

To quantify the relation between ccQTLs and GWAS variants, we calculated the λ-inflations. Specifically, we used the fine-mapped variants from the disease GWAS as taken from the Open Targets fine-mapping, and calculated the λ-inflation^66^ for the ccQTL variants that overlap with GWAS variants.

### Annotation of GWAS-linked independent loci

To assess which features of our independent sc-eQTLs are shared with loci associated to GWAS or associated to bulk *cis*-eQTLs from eQTLGen, we leveraged two logistic regression models. One predicting independent loci with disease GWAS colocalization (yes = 1, no = 0), and another one predicting loci that are shared with bulk *cis*-eQTLs from eQTLGen effect (yes = 1, no = 0). As predictors we included: the disease EFO category, and the number of cell types and genes an independent locus is identified in; whether the locus is shared among the two major linages (lineage mixing) and shared with bulk *cis*-eQTLs from eQTLGen. The logistic regression models were fitted with the glm function in R 4.2.1.

### Variant-based GWAS enrichment of eQTLs

All fine-mapped eQTL loci were used for variant-based GWAS enrichment. We annotated these variants with GWAS variants with genome-wide significant (P < 5 x 10^−8^) associations (v1.0.2) downloaded from the GWAS Catalog, while considering LD. We used EFO terms to group variants into different trait groups. We mapped the GWAS traits to EFO-IDs^10^ and corresponding parent terms from the GWAS catalog^67^ using gwasrapidd v0.99.17. The different trait categories were defined as follow:

- All traits: no filtering; all traits in GWAS catalog
- Disease GWAS: only kept traits assigned to these EFO parent terms: ‘Cancer (EFO_0000616)’, ‘Immune system disorder (EFO_0000540)’, ‘Other disease (EFO_0000408)’, ‘Cardiovascular disease (EFO_0000319)’, ‘Neorological disorder (EFO_0000618)’, ‘Metabolic disorder (‘EFO_0000589’)
- Immune disease GWAS: only kept traits assigned to these EFO parent terms:’Immune system disorder (EFO_0000540)’

To analyze the enrichment across all cell types (‘All’ category) we used only one cell type observation per eGene. If an eGene had an eQTL effect in multiple cell types, the observation of the cell type with the lowest p-value of the eQTL effect was used.

In sample LD information was used to filter the eQTL and complete set of assessed variants (R2 ≥ 0.1) as well as determining whether the GWAS variants are in LD (R2 ≥ 0.8). The eSNPs were filtered such that the most significant variant in high LD was kept. eSNPs are determined to be GWAS variants either if they are present in the GWAS catalog set of GWAS variants or if they are in LD with any of the GWAS variants.

### eGene enrichment analysis

To gain insight into the potential upstream regulator that may be associated to the *trans*-eGenes that were linked to the same *cis*-eGenes, we performed gene set enrichments using the R package g:profiler^68^. We tested only *trans*-eGenes that formed *cis*-*trans* gene-gene pairs in which the same *cis-*eGene was linked to at least 9 other *trans*-eGenes. All settings were left at defaults provided by the tool developers, other than the background sets. The background gene set consisted of all genes that showed at least one *trans*-eQTLs in eQTLGen.

Additionally, to get higher resolution on the relevance of TFs we prepared specific test information by parsing the information from the ReMap project^69^. Specifically, we added three extra information sources, we added the *cis*-regulatory modules (CRMs) as defined by ReMap, the non-redundant TF information over all cell types and the non-redundant TF information for blood relevant cell types^70^. These information sources were added by providing custom data to g:profiler^68^.

## Code availability

The pipelines for (WG1) genotype imputation and scRNA-seq data preprocessing, QC and donor demultiplexing; (WG2) cell type assignment; (WG3) eQTL mapping and meta-analysis and downstream analyses, can be found on Github (https://github.com/sc-eQTLgen-consortium). The pipelines are established in Snakemake^48^, and all software is provided in a Singularity^71^ or Docker^72^ image for consistency across datasets.

## Data availability

Full eQTL summary statistics will be made available after publication the sc-eQTLGen consortium webpage: https://www.eqtlgen.org/sc/. eQTLGen summary statistics will be publicly available upon acceptance of the respective paper.

## Ethical compliance

All cohorts included in this study enrolled participants with informed consent and collected and analyzed data in accordance with ethical and institutional regulations. The information about individual institutional review board approvals is available in the original publications for each cohort. All procedures performed in studies involving human participants were in accordance with the ethical standards of the institutional and/or national research committee and with the 1964 Helsinki declaration and its later amendments or comparable ethical standards.

## Acknowledgment

We thank all the children who participated in the Burkina Faso study and their families. We thank the staff of the Centre National de Recherche et de Formation sur le Paludisme and of the Groupe de Recherche Action en Santé for facilitating sample collection and clinical work. We acknowledge support from the National Institute for Health Research (NIHR) Cambridge Biomedical Research Centre, United Kingdom. The images in Figure 4f are created using Servier Medical Art (https://smart.servier.com), licensed under CC BY 4.0. We thank Katherine McIntyre for the English editing of our manuscript. The authors wish to acknowledge the contributions of all participating cohorts and study participants.

## Author contributions

Conceived and designed the computational framework: D.K., L.M., D.N., A.R., O.S., M.Hemberg, A.M., M.M., J.P., M.J.B., L.F., M.W.

Analyzed the data: D.K., L.M., D.N., A.R., J.A., M.K., J.T.H.L., R.O., M.V., R.W., Y.A., M.Ban, O.B., M.Berg, I.B., M.D., R.E., M.G.G., H.G., M.Heinig, P.H., C.H., Y.I., P.K., N.K., W.L., Y.L., C.L., V.M., J.M., M.C.N., M.N., J.N., Y.O., S.S., I.S., W.P., D.R., T.S., J.S., G.T., U.V., S.Y., J.Y., Z.Z., eQTLGen consortium, M.J.B., M.W.

Performed statistical analysis: D.K., L.M., D.N., A.R., D.C., M.Heinig, C.L., Y.T., G.T.

Contributed analysis tools: D.K., L.M., D.N., A.R., J.A., M.K., J.T.H.L., R.O., M.V., R.W., D.C., Y.T., G.T.

Contributed dataset: D.K., M.K., R.O., Y.A., M.Ban, O.B., M.Berg, I.B., M.D., R.E., M.G.G., H.G., P.H., C.H., Y.I., P.K., N.K., W.L., Y.L., V.M., J.M., H.N., Y.M., M.C.N., M.N., J.N., Y.O., S.S., I.S., W.P., D.R., T.S., J.S., G.T., U.V., S.Y., J.Y., Z.Z., eQTLGen, J.P. M.J.B., L.F., M.W.

Jointly supervised research: M.J.B., L.F., M.W. Wrote the paper: M.J.B., M.W.

## Figures

## Tables

## Supplementary Figures

## Supplementary Tables

