## Supplementary Notes for "Federated single-cell QTL meta-analysis reveals novel disease mechanisms"

**Supplementary Figures**

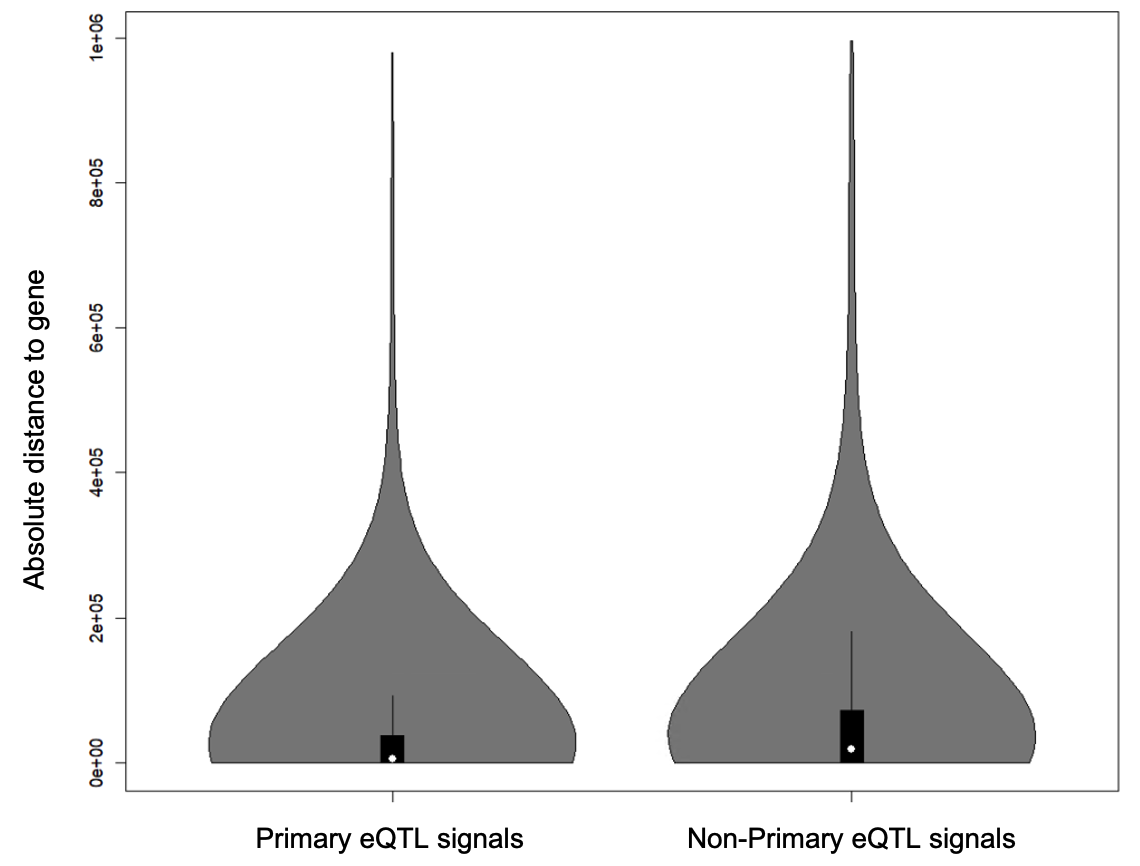

**Figure S1. Location distribution of primary and non-primary eQTL signals relative to the gene location, depicting absolute distance.**

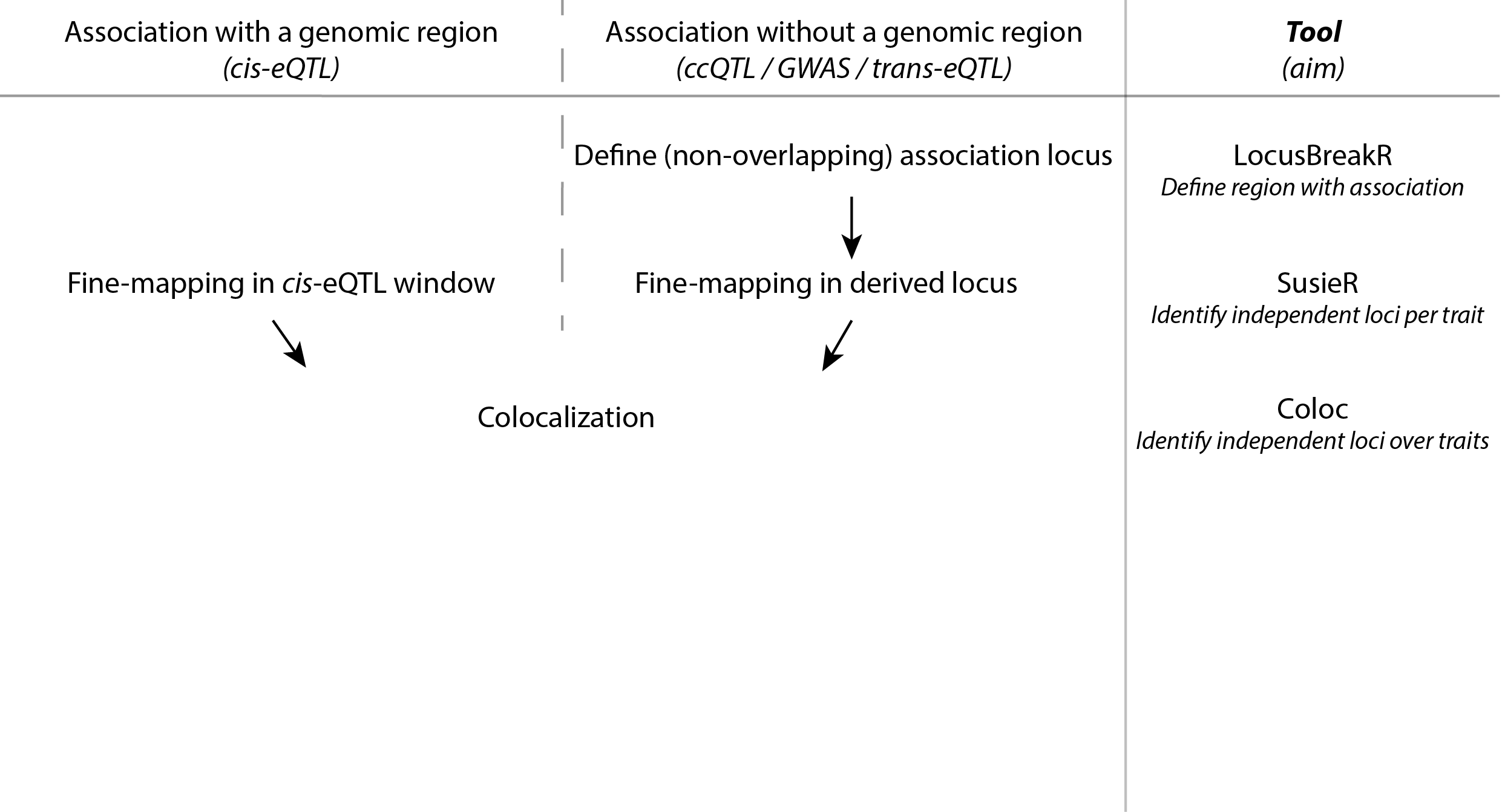

**Figure S2. The fine-mapping and colocalization approach for QTLs and GWAS traits.**

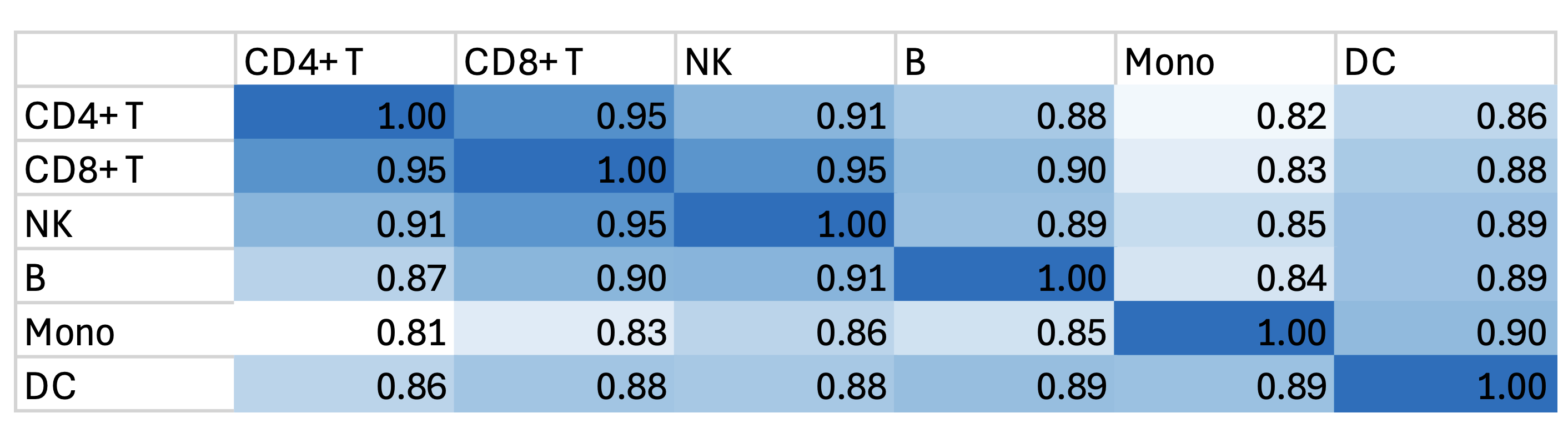

**Figure S3. Fine-mapping- and colocalization-based replication of eQTL effects across cell types.**

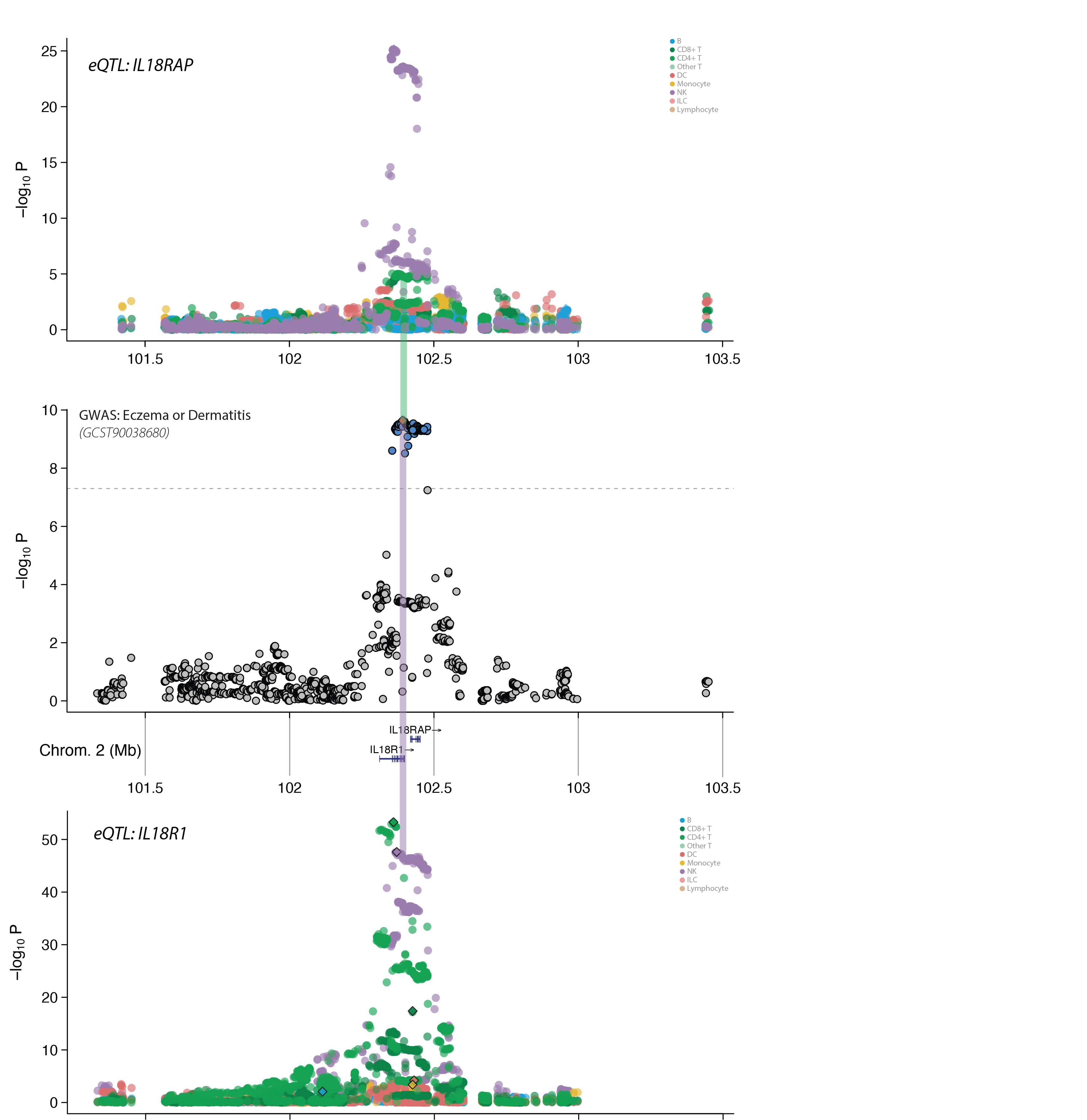

**Figure S4. Colocalization of single-cell eQTLs in the *IL18R1* / *IL18RAP* locus to the GWAS on ‘Eczema or Dermatitis’.** Locus plot for the colocalization between the GWAS signal (rs12712145, ‘GCST90038680: Eczema or dermatitis’) (center) and *IL18RAP* in CD4+ T cells (top), and *IL18R1* in NK cells (bottom).

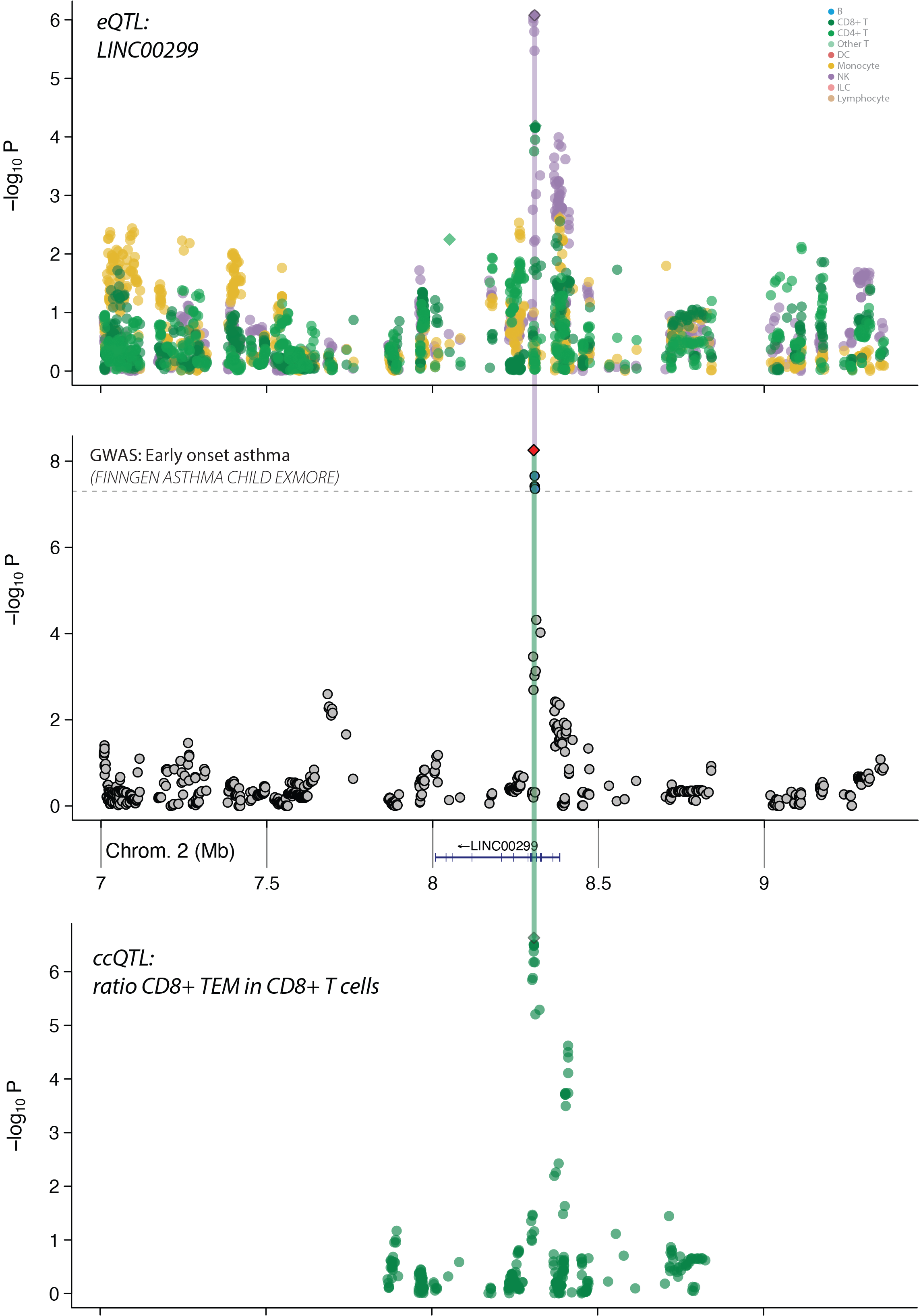

**Figure S5. Shared eQTL and ccQTL effect in the Early onset asthma locus (rs391934).**

Locus plot for the colocalization between the GWAS signal (rs391934, “FINNGEN_R12_ASTHMA_CHILD_ EXMORE: Childhood Asthma’) (center) and *LINC00299* in NK cells (top). The same eQTL also colocalizes with the ccQTL that changes the relative proportion of CD8+ TEM cells within CD8+ T cells (bottom).

**Supplementary Tables**

Table S1. Donor characteristics per cohort.

Table S2. Number of cells passed QC per cohort and analysis.

Table S3. Cell count QTLs.

Table S4. Expression QTLs.

Table S5. GWAS colocalization of eQTLs.

Table S6. LD-based GWAS enrichment of eQTLs. **A.** enrichment of eQTL variants among GWAS variants relative to tested variants within sc-eQTLGen. Enrichments shown are for all GWAS, disease GWAS and Immune related GWAS variants. Shown for the cell type-specific eQTL summary statistics, the combined single-cell eQTL summary statistics and bulk eQTLGen summary statistics. **B.** Z-test comparing eQTL enrichments for bulk-based eQTL summary statistics versus single cell eQTL summary statistics.

Table S7. Colocalization of single-cell *cis*-eQTLs with bulk *trans*-eQTLs.

**Supplementary notes**

**Note S1: eQTLGen Consortium – Author information**

Author list is ordered alphabetically.

Habibul Ahsan^1^, Marta E. Alarcón-Riquelme^2,3^, Philip Awadalla^4^, Guillermo Barturen^5,6,2^, Alexis Battle^7,8,9^, Frank Beutner^10^, Cornelis Blauwendraat^11,12^, Collins Boahen^13,14^, Toni Boltz^15^, Dorret I. Boomsma^16,17,18^, Andrew Brown^19^, John Budde^20,21^, Katie L. Burnham^22^, John Chambers^23,24,25^, Evans Cheruiyot^26^, Surya B. Chhetri^7,27^, Annique Claringbould^28^, IMI DIRECT Consortium^29^, PRECISESADS Clinical Consortium^2^, Carlos Cruchaga^20,21,30,31,32,33^, Kensuke Daida^11,12,34^, Emma E. Davenport^22^, Théo Dupuis^19^, Diptavo Dutta^35^, Tõnu Esko^36^, Aiman Farzeen^37,38,39^, Marie-Julie Favé^40,4^, Luigi Ferrucci^41^, Lude H. Franke^42,43^, Timothy M. Frayling^44^, Koichi Fukunaga^45^, J. Raphael Gibbs^46^, Greg Gibson^47^, Christian Gieger^38,39^, Marleen van Greevenbroek^48,49^, Binisha Hamal Mishra^50,51,52^, Takanori Hasegawa^53^, Jouke Jan Hottenga^54,55^, M. Arfan Ikram^56^, Michael Inouye^57,58,59,60,61,62^, Rick Jansen^63^, Farzana Jasmine^64^, Matt Johnson^20,21^, Mika Kähönen^51,65^, Carla J.H van der Kallen^48,49^, Muhammad G. Kibriya^64^, Holger Kirsten^66,67^, Julian C. Knight^68^, Knut Krohn^69^, Viktorija Kukushkina^36^, Vinod Kumar^13,14,42,70^, Sandra Lapinska^71^, Terho Lehtimäki^50,51,52^, Yun Li^72,73,74^, Markus Loeffler^67,66^, Marie Loh^23,75,76,25^, Leo-Pekka Lyytikäinen^50,51,52^, Reedik Mägi^36^, Javier Martin^77^, Angel Martinez-Perez^78,79^, Allan F. McRae^26^, Joyce van Meurs^80,81^, Pashupati P. Mishra^50,51,52^, Younes Mokrab^82,83,84^, Grant W. Montgomery^26^, Juha Mykkänen^85,86^, Haroon Naeem^82^, Sini Nagpal^47^, Ho Namkoong^87^, Matthias Nauck^88^, Yukinori Okada^89,90,91,92,93^, Roel Ophoff^71,15,94^, Katja Pahkala^86,85,95^, Bogdan Pasaniuc^96^, Dirk S. Paul^57,58,97^, Brenda W.J.H Penninx^63^, Elodie Persyn^57,58,59^, Annette Peters^39,98,99,100^, Brandon Pierce^64^, René Pool^55,17^, Holger Prokisch^101,37,102^, Laura M. Raffield^73^, Venket Raghavan^67^, Olli T. Raitakari^86,103,85,104^, Emma Raitoharju^105,106^, María Rivas-Torrubia^2^, Ruth D. Rodríguez^2^, Suvi P. Rovio^86,85,107^, Jessie Sanford^20,21^, Casper G. Schalkwijk^48,49^, Markus Scholz^67,66^, Andrew Singleton^108^, Eline Slagboom^109^, José Manuel Soria^78,79^, Juan Carlos Souto^78^, Yun Ju Sung^20,21^, Darwin Tay^23^, Estonian Biobank Research Team^36^, The HELIOS Study Team^23^, Alexander Teumer^110,111,112^, Joachim Thiery^66,113^, Alex Tokolyi^22^, Lin Tong^64^, Jan Veldink^114^, Joost Verlouw^80^, Ana Viñuela^115,19^, Peter M. Visscher^26^, Uwe Völker^116,111^, Urmo Võsa^36^, Qingbo S. Wang^89,90,91^, C.A. Robert Warmerdam^42,43^, Stefan Weiss^116,111^, Jia Wen^73^, Harm-Jan Westra^42,43^, Andrew R. Wood^117^, Manke Xie^47^

1. Biological Sciences Division, Institute for Population and Precision Health (IPPH), University of Chicago, Chicago, IL, USA

2. Pfizer–University of Granada–Junta de Andalucía Centre for Genomics and Oncological Research, Granada, Spain

3. Institute of Environmental Medicine, Karolinska Institute, Stockholm, Sweden

4. Ontario Institute for Cancer Research, Toronto, Ontario, Canada

5. Department of Genetics, Faculty of Science, University of Granada, 18071 Granada, Spain

6. Bioinformatics Laboratory, Biotechnology Institute, Centro de Investigación Biomédica, PTS, Avda. del Conocimiento s/n, 18100 Granada, Spain

7. Department of Biomedical Engineering, Johns Hopkins University, Baltimore, MD, USA

8. Department of Computer Science, Johns Hopkins University, Baltimore, MD, USA

9. Department of Genetic Medicine, Johns Hopkins University, Baltimore, MD, USA

10. Department of Internal Medicine/Cardiology, Heart Center Leipzig at Leipzig University, Leipzig, Germany

11. Integrative Neurogenomics Unit, Laboratory of Neurogenetics, National Institute on Aging, National Institutes of Health, Bethesda, MD, USA

12. Center for Alzheimer’s and Related Dementias (CARD), National Institute on Aging and National Institute of Neurological Disorders and Stroke, National Institutes of Health, Bethesda, MD, USA

13. Department of Internal Medicine and Radboud Institute of Molecular Life Sciences (RIMLS), Radboud University Medical Center, Nijmegen, 6525 HP, the Netherlands

14. Department of Internal Medicine and Radboud Center for Infectious Diseases (RCI), Radboud University Medical Center, Nijmegen, 6525 HP, the Netherlands

15. Department of Human Genetics, David Geffen School of Medicine, University of California Los Angeles, Los Angeles, CA, USA

16. Department of Complex Trait Genetics, Center for Neurogenomics and Cognitive Research, Vrije Universiteit Amsterdam, the Netherlands

17. Amsterdam Public Health research institute, Amsterdam, the Netherlands

18. Amsterdam Reproduction and Development (AR&D) Research Institute, Amsterdam, the Netherlands

19. Population Health and Genomics, University of Dundee, Dundee, Scotland, UK

20. Department of Psychiatry, Washington University School of Medicine, St. Louis, MO, United States

21. NeuroGenomics and Informatics Center, Washington University School of Medicine, St. Louis, MO 63110, USA

22. Wellcome Sanger Institute, Wellcome Genome Campus, Hinxton, UK

23. Lee Kong Chian School of Medicine, Nanyang Technological University, Singapore

24. Precision Health Research (PRECISE), Singapore

25. Department of Epidemiology and Biostatistics, School of Public Health, Imperial College London, United Kingdom

26. Institute for Molecular Bioscience, The University of Queensland, Brisbane, Australia

27. Center for Computational Biology, Johns Hopkins University, Baltimore, MD, USA

28. Department of Internal Medicine, Erasmus MC, Erasmus University Medical Center Rotterdam, Rotterdam, the Netherlands

29. https://directdiabetes.org/

30. Department of Neurology, Washington University School of Medicine, St. Louis, MO 63110, USA

31. Knight Alzheimer Disease Research Center, Washington University School of Medicine, St. Louis, MO, United States

32. Hope Center for Neurological Disorders, Washington University School of Medicine, St. Louis, MO, United States

33. Dominantly Inherited Alzheimer Disease Network (DIAN)

34. Department of Neurology, Faculty of Medicine, Juntendo University, Tokyo, Japan

35. Division of Cancer Epidemiology & Genetics, National Cancer Institute, Bethesda, MD, USA

36. Estonian Genome Centre, Institute of Genomics, University of Tartu, Tartu, Estonia

37. Institute of Neurogenomics, Computational Health Center, Helmholtz Munich, Neuherberg, Germany

38. Research Unit of Molecular Epidemiology, Helmholtz Zentrum München - German Research Center for Environmental Health, Neuherberg, Germany

39. Institute of Epidemiology, Helmholtz Zentrum München - German Research Center for Environmental Health, Neuherberg, Germany

40. Concordia University, Centre for Functional and Structural Genomics, Biology department, Montreal, Canada

41. Translational Gerontology Branch, National Institute on Aging, National Institutes of Health, Baltimore, MD, USA

42. Department of Genetics, University Medical Center Groningen, University of Groningen, Groningen, the Netherlands

43. Oncode Institute, Utrecht, the Netherlands

44. Department of Genetic Medicine and Development, CMU, University of Geneva

45. Division of Pulmonary Medicine, Department of Medicine, Keio University School of Medicine, Tokyo, Japan

46. Computational Biology Group, Laboratory of Neurogenetics, National Institute on Aging, Bethesda, MD, USA

47. Center for Integrative Genomics, Georgia Institute of Technology, Atlanta, GA, USA.

48. Cardiovascular Research Institute Maastricht (CARIM), Maastricht University, Maastricht, the Netherlands

49. Department of Internal Medicine, Maastricht University Medical Center, Maastricht, the Netherlands

50. Department of Clinical Chemistry, Faculty of Medicine and Health Technology, Tampere University, Tampere, Finland

51. Finnish Cardiovascular Research Center Tampere, Faculty of Medicine and Health Technology, Tampere University, Tampere, Finland

52. Department of Clinical Chemistry, Fimlab Laboratories, Tampere, Finland

53. M&D Data Science Center, Tokyo Medical and Dental University, Tokyo, Japan

54. Neurological Disorder Research Center, Qatar Biomedical Research Institute (QBRI), Hamad Bin Khalifa University (HBKU), Qatar Foundation, Doha P.O. Box 5825, Qatar

55. Department of Biological Psychology, Vrije Universiteit Amsterdam, Amsterdam, the Netherlands

56. Department of Epidemiology, Erasmus MC University Medical Center, Rotterdam, the Netherlands

57. British Heart Foundation Cardiovascular Epidemiology Unit, Department of Public Health and Primary Care, University of Cambridge, Cambridge, UK

58. Victor Phillip Dahdaleh Heart and Lung Research Institute, University of Cambridge, Cambridge, UK

59. Cambridge Baker Systems Genomics Initiative, Department of Public Health and Primary Care, University of Cambridge, Cambridge, UK

60. Cambridge Baker Systems Genomics Initiative, Baker Heart and Diabetes Institute, Melbourne, VIC, Australia

61. Health Data Research UK Cambridge, Wellcome Genome Campus and University of Cambridge, Cambridge, UK

62. British Heart Foundation Centre of Research Excellence, University of Cambridge, Cambridge, UK

63. Department of Psychiatry, Amsterdam Public Health (Mental Health program) and Amsterdam Neuroscience (Mood, Anxiety, Psychosis, Stress and Sleep Program) Research Institutes, Amsterdam UMC Location Vrije University Amsterdam, Amsterdam, Netherlands

64. Biological Sciences Division, Public Health Sciences, University of Chicago, Chicago, IL, USA

65. Department of Clinical Physiology, Tampere University Hospital, Tampere Finland

66. LIFE – Leipzig Research Center for Civilization Diseases, Leipzig University, Leipzig, Germany

67. Institute for Medical Informatics, Statistics and Epidemiology, Leipzig University, Leipzig, Germany

68. Centre for Human Genetics, University of Oxford, Oxford, UK

69. Medical Faculty, University of Leipzig, Leipzig, Germany

70. Nitte (Deemed to Be University), Medical Sciences Complex, Nitte University Centre for Science Education and Research (NUCSER), Deralakatte, Mangalore, 575018, India

71. Bioinformatics Interdepartmental Program, University of California Los Angeles, Los Angeles, CA, USA

72. Department of Biostatistics, University of North Carolina at Chapel Hill, Chapel Hill, NC, USA

73. Department of Genetics, University of North Carolina, Chapel Hill, NC, USA

74. Department of Computer Science, University of North Carolina at Chapel Hill, Chapel Hill, NC, USA

75. National Skin Centre, Research Division, Singapore

76. Genome Institute of Singapore, Agency for Science, Technology and Research, Singapore

77. Instituto de Parasitología y Biomedicina López-Neyra, Consejo Superior de Investigaciones Científicas (IPBLN-CSIC), Granada, Spain

78. Unit of Genomics of Complex Diseases, Institut de Recerca Sant Pau (IR Sant Pau), Sant Quintí 77-79, 08041 Barcelona, Spain

79. Centre for Biomedical Network Research on Rare Diseases (CIBERER), Instituto de Salud Carlos III, Madrid, Spain

80. Department of Internal Medicine, Erasmus MC University Medical Center, Rotterdam, the Netherlands

81. Department of Orthopaedics and Sportsmedicine, Erasmus MC University Medical Center, Rotterdam, the Netherlands

82. Sidra Medicine, Doha, Qatar

83. Department of Biomedical Science, College of Health Sciences, Qatar University, Doha, Qatar

84. Department of Genetic Medicine, Weill Cornell Medicine, Qatar

85. Centre for Population Health Research, University of Turku and Turku University Hospital, Turku, Finland

86. Research Centre of Applied and Preventive Cardiovascular Medicine, University of Turku, Turku, Finland

87. Department of Infectious Diseases, Keio University School of Medicine, Tokyo, Japan

88. Institute of Clinical Chemistry and Laboratory Medicine, University Medicine Greifswald, Greifswald, Germany

89. Department of Genome Informatics, Graduate School of Medicine, the University of Tokyo, Tokyo, Japan

90. Department of Statistical Genetics, Osaka University Graduate School of Medicine, Suita, Japan

91. Laboratory for Systems Genetics, RIKEN Center for Integrative Medical Sciences, Yokohama, Japan

92. Laboratory of Statistical Immunology, Immunology Frontier Research Center (WPI-IFReC), Osaka University, Suita, Japan

93. Premium Research Institute for Human Metaverse Medicine (WPI-PRIMe), Osaka University, Suita, Japan

94. Center for Neurobehavioral Genetics, Semel Institute for Neuroscience and Human Behavior, David Geffen School of Medicine, University of California Los Angeles, Los Angeles, USA

95. Paavo Nurmi Centre & Sports and Exercise Medicine, University of Turku, Turku, Finland

96. Department of Genetics, Perelman School of Medicine, University of Pennsylvania, Philadelphia, PA, USA

97. Centre for Genomics Research, Discovery Sciences, BioPharmaceuticals R&D, AstraZeneca, Cambridge, UK

98. Chair of Epidemiology, IBE, Faculty of Medicine, LMU Munich, Munich, Germany

99. German Centre for Cardiovascular Research (DZHK), Partner Site Munich Heart Alliance, Munich, Germany

100. German Center for Diabetes Research (DZD), Neuherberg, Germany

101. School of Medicine, Institute of Human Genetics, Technical University of Munich, Munich, Germany

102. German Center for Child and Adolescent Health (DZKJ), partner site Munich, Munich, Germany

103. Department of Clinical Physiology and Nuclear Medicine, Turku University Hospital, Turku, Finland

104. InFLAMES Research Flagship, University of Turku, Turku, Finland

105. Molecular Epidemiology, Faculty of Medicine and Health Technology, Tampere University, Tampere, Finland

106. Tampere University Hospital, Tampere, Finland

107. Department of Public Health, University of Turku and Turku University Hospital, Turku, Finland

108. Laboratory of Neurogenetics, National Institute on Aging, National Institutes of Health, Bethesda, MD, USA

109. Section of Molecular Epidemiology, Department of Biomedical Data Sciences, Leiden University Medical Center, Leiden, the Netherlands

110. Department of Psychiatry and Psychotherapy, University Medicine Greifswald, Greifswald, Germany

111. DZHK (German Center for Cardiovascular Research), Partner Site Greifswald, Greifswald, Germany

112. Department of Population Medicine and Lifestyle Diseases Prevention, Medical University of Bialystok, Bialystok, Poland

113. Institute of Laboratory Medicine, Clinical Chemistry and Molecular Diagnostics, University of Leipzig Medical Center, Leipzig, Germany

114. Department of Neurology, UMC Utrecht Brain Center Rudolf Magnus, Utrecht, the Netherlands

115. Biosciences Institute, Faculty of Medical Sciences, University of Newcastle, Newcastle upon Tyne, UK

116. Interfaculty Institute of Genetics and Functional Genomics, University Medicine Greifswald, Greifswald, 17475, Germany

117. College of Medicine and Health, University of Exeter, Exeter, UK

**Note S2: Cohort descriptions and cohort specific methods.**

**Trynka cohort**

The Trynka cohort includes 119 healthy adults from British ancestry^1^. Blood was collected in EDTA tubes, and PBMCs were isolated using Ficoll-Paque PLUS density gradient centrifugation. Genotyping of PBMC genomic DNA was performed using the Infinium CoreExome-24 (v1.3) chip. Naïve (CD25−, CD45RA+) and memory (CD25−, CD45RO+) CD4+ T cells were isolated from PBMCs using the EasySep naïve CD4+ T cell isolation kit and memory CD4+ T cell enrichment kit (StemCell Technologies), according to the manufacturer’s instructions. T cells were then stimulated with anti-CD3/anti-CD28 human T-Activator Dynabeads® (Invitrogen) at a 1:2 ratio of beads to T cells, or left unstimulated. Cells were harvested after 16 h, 40 h, and 5 d of stimulation, or left unstimulated for 16h. Next, CD4+ T cells were stained with a live/death dye (DAPI), and dead cells were removed using FACS. The unstimulated live CD4+ T cells were used for scRNA-seq using 10X Genomics 3’-end v2 chemistry reagents on sample pools of 4−6 donors each. Libraries were sequenced using a custom program (26-8-0-98) on an Illumina HiSeq4000 using a 75-bp paired-end kit. Cell Ranger v3.0.0 software was used for basic alignment and initial processing of the scRNA-seq data. The genotypes and raw scRNA-seq data have been deposited at the European Genome-Phenome Archive (EGA) under accession numbers [EGAD00001008197](https://ega-archive.org/datasets/EGAD00001008197), [EGAD00010002291](https://ega-archive.org/datasets/EGAD00010002291) and [EGAD00001008197](https://ega-archive.org/datasets/EGAD00001008197), respectively. After QC (**Methods section: WP1-3**), 117 individuals remained for the eQTL meta-analysis.

**Wijst 2018 cohort**

The Wijst 2018 dataset includes 45 adults from the general population of the northern part of the Netherlands^2^, part of the Lifelines-DEEP cohort^3^. Blood was collected in EDTA tubes, and PBMCs were isolated using sodium-heparin CPT tubes. PBMCs were cryopreserved until genotyping and scRNA-seq. Genotyping was performed using the HumanCytoSNP-12 BeadChip. scRNA-seq was conducted using 10X Genomics 3’-end v2 chemistry reagents on sample pools of 5−6 donors each. Libraries were sequenced using a custom program (27-9-0-138) on an Illumina HiSeq4000 using a 75-bp paired-end kit. Cell Ranger v1.3 software was used for basic alignment and initial processing of the scRNA-seq data. The processed (de-anonymized) scRNA-seq data, including a text file that links each cell barcode to its respective individual, have been deposited at the EGA under accession number [EGAS00001002560](https://www.ebi.ac.uk/ega/studies/EGAS00001002560). After QC (**Methods section: WP1-3**), 40 individuals remained for the eQTL meta-analysis.

**Li 300-BCG cohort**

The Li-BCG study contains 39 healthy adults of Western European ancestry^4^, part of the 300-BCG cohort set up to study the immunological effects of BCG vaccination^5^**.** These 39 individuals were chosen from the larger 300-BCG cohort (N = 325 individuals) based on the strength of their trained immunity response, as measured by a change in IL-1β response upon 24h 5×10^6^ CFU/mL heat-killed *S. aureus* exposure before and 3 months after 0.1 ml *in vivo* BCG vaccination (BCG-Bulgaria, InterVax): high (FC ≥ 2 of *S. aureus*-induced IL-1β production, N = 19) and low (FC < 2, N = 20) responders. Blood was collected in EDTA tubes (from before and 3 months after 0.1 ml BCG vaccination), and PBMCs were isolated using density centrifugation over Ficoll-Paque. PBMCs were cryopreserved until genotyping and scRNA-seq. Genotyping was done using the Infinium Global Screen Array (GSA) MD v1.0. PBMCs, either left untreated in RPMI medium or exposed for 4h to 10 ng/ml LPS (serotype 055:B5, Sigma), were used for scRNA-seq. scRNA-seq was conducted using 10X Genomics 3’-end v3.1 chemistry reagents on sample pools of 3−5 donors per library. Libraries were sequenced using a custom program (28-8-90) on an Illumina NovaSeq6000 using a 100-bp paired-end kit. Cell Ranger v3.1.0 software was used for basic alignment and initial processing of the scRNA-seq data. The processed (de-anonymized) scRNA-seq data, including a text file that links each cell barcode to its respective individual, have been deposited at the EGA under accession number [EGAD00001010055](https://ega-archive.org/datasets/EGAD00001010055). For the eQTL meta-analysis, only the scRNA-seq data from the timepoint before BCG vaccination (RPMI medium control condition) was used.

**OASIS cohort**

The OASIS (Osaka Atlas of Immune Cells) dataset includes 96 healthy adults from the general Japanese population^6–8^. Blood was collected from these individuals in heparin tubes, and PBMCs were isolated using Leucosep density gradient centrifugation. PBMCs were cryopreserved until genotyping and scRNA-seq. Genotyping was performed using the Infinium Asian Screening Array. scRNA-seq was conducted using 10X Genomics 5’-end v1.1 or v2 chemistry reagents, each library containing a single individual. Libraries were sequenced using a custom program (v1.1: 26-8-0-91, v2 26-10-10-90) on an Illumina NovaSeq 6000 using a 100-bp paired-end kit. Cell Ranger v5.0.0 software was used for basic alignment and initial processing of the scRNA-seq data. The processed (de-anonymized) scRNA-seq data have been deposited at the Japanese Genotype-Phenotype Archive (JGA) under accession numbers [JGAS000593](https://ddbj.nig.ac.jp/resource/jga-study/JGAS000593) and [JGAD000722](https://ddbj.nig.ac.jp/resource/jga-dataset/JGAD000722). The genotype data have been deposited at the EGA under accession number [EGAS00001006950](https://ega-archive.org/studies/EGAS00001006950). After QC (**Methods section: WP1-3**), 91 individuals remained for the eQTL meta-analysis.

**Ye (SLE) cohort -** **CLUES and ImmVar cohorts**
The Ye SLE dataset^9^ includes 162 female systematic lupus erythematosus patients from donors in the California Lupus Epidemiological Study (CLUES) cohort, 49 matching healthy controls from the UCSF Rheumatology Clinic, along with 46 additional healthy samples from the Immune Variation Project (ImmVar), including donors of East Asian and European descent. Blood was collected in EDTA tubes, and PBMCs were isolated using density centrifugation over Ficoll-Paque. PBMCs were cryopreserved until genotyping and scRNA-seq. Genotyping was performed using the Affymetrix World LAT array (CLUES and UCSF Rheumatology Clinic cohort) and the OmniExpressExome chip (ImmVar cohort). Due to the different chips used for genotyping, the ImmVar cohort was processed separately through the sc-eQTLgen pipelines. scRNA-seq was conducted using 10X Genomics 3’-end v2 chemistry reagents on sample pools of 16 donors each. Libraries were sequenced using a custom program (26-8-0-98) on an Illumina HiSeq4000 or NovaSeq6000 using a 100-bp paired-end kit. Cell Ranger v2.0 software was used for basic alignment and initial processing of the scRNA-seq data. The genotype data is available on dbGap ([phs002812.v1.p1](https://www.ncbi.nlm.nih.gov/projects/gap/cgi-bin/study.cgi?study_id=phs002812.v1.p1)). The processed and demultiplexed scRNA-seq data have been deposited at the Gene Expression Omnibus (GEO) under accession number [GSE174188](https://www.ncbi.nlm.nih.gov/geo/query/acc.cgi?acc=GSE174188). After QC (**Methods section: WP1-3**), 170 individuals remained for the eQTL meta-analysis.

**Powell OneK1K cohort**

The OneK1K study^10^ includes 1,029 participants from the Royal Hobart Hospital, Hobart Eye Surgeons, or retirement villages in Hobart (Australia) without immune disease. All participants were Northern European ancestries, with the majority (73%) being 60 years or older. Blood was collected, and PBMCs were isolated using sodium-heparin CPT tubes. Live cells were isolated with the Miltenyi Dead Cell Removal Kit. PBMCs were cryopreserved until genotyping and scRNA-seq. Genotyping was performed using the Infinium GSA. For single-cell experiments, PBMCs from 12−14 participants were pooled and assayed with 10x Genomics 3’-end v2. Libraries were sequenced using a custom program (26-8-0-98) on an Illumina Novaseq6000 using a 100-bp paired-end kit. Cell Ranger v2.2.0 software was used for alignment and processing. The genotype and scRNA-seq data are available on the GEO under accession number [GSE196830](https://www.ncbi.nlm.nih.gov/geo/query/acc.cgi?acc=GSE196830). Following QC (**Methods section: WP1-3**), 1,017 individuals remained for eQTL detection.

**Asthma Remission (ARMS) cohort**

The ARMS cohort^11^ (unpublished) includes 37 adults who had childhood onset asthma, as confirmed by retrospective inspection of the original lung function results used for diagnosis, and 13 healthy controls. The (ex-)asthma patients have either active disease (persistent asthma, N = 14) or are in clinical (N = 13) or complete remission (N = 10), as evaluated during the clinical phenotyping of these patients. Patients with asthma stopped using inhaled corticosteroid 6 weeks before blood was drawn. Whole blood was collected in lithium heparin blood tubes. Genotyping was performed using the Infinium GSA-24 v3.0. For scRNA-seq, blood was further processed by lysing the red blood cells using an ammonium chloride-potassium solution (155 mM ammonium chloride (NH_4_Cl), 10 mM potassium bicarbonate (KHCO_3_), 0.1 mM EDTA). Cells were centrifuged for 5 min at 4°C, 550g, after which the cell pellet was washed twice with PBS containing 1% BSA, followed by staining for cell surface markers (CD3, CD4) and propidium iodide (live/dead). FACS sorting was used for selecting live, CD3 and CD4 double positive cells (CD4+ T cells). These CD4+ T cells were sorted into 96-well plates for SmartSeq2 scRNA-seq using a MoFlo Astrios with Summit Software. Libraries were sequenced on an Illumina HiSeq 4000. The processed (de-anonymized) scRNA-seq data have been deposited at the EGA under accession number [EGAD00001005064](https://ega-archive.org/datasets/EGAD00001005064). After QC (**Methods section: WP1-3**), 45 individuals remained for the eQTL meta-analysis.

**Oelen 1M-scBloodNL cohort**

The Oelen cohort includes 120 individuals residing in the northern part of the Netherlands^12^. These participants are also part of the LifeLines-DEEP cohort^3^. From these individuals, EDTA blood was drawn and PBMCs were isolated using sodium-heparin CPT tubes. PBMCs were cryopreserved until genotyping and scRNA-seq. Genotyping was conducted using the HumanCytoSNP-12 v2.1 array. PBMCs, either left untreated or exposed for either 3h or 24h to heat-killed *C. albicans blastoconidia*, *M. Tuberculosis*, or *P. Aeruginosa*, were used for scRNA-seq. scRNA-seq was conducted using 10X Genomics 3’ scRNA-seq v2 (80 donors) and v3 chemistry (40 donors) reagents on sample pools of 8 donors each. Libraries were sequenced using a custom program (27-9-150 (v2), 28-8-150 (v3)) on an Illumina NovaSeq6000 using a 100-bp paired-end kit. Cell Ranger v3.0.2 software was used for basic alignment and initial processing of the scRNA-seq data. The processed (de-anonymized) scRNA-seq data, including a text file that links each cell barcode to its respective individual, have been deposited at the EGA under accession number [EGAS00001005376](https://www.eqtlgen.org/sc/datasets/1m-scbloodnl-dataset.html). For this meta-analysis, only the scRNA-seq data from the untreated cells was used. After QC (**Methods section: WP1-3**), 107 (v2: 74 donors, v3: 33 donors) individuals remained for the eQTL meta-analysis.

**Van Blokland (STEMI) cohort**

The Van Blokland cohort includes 38 adult patients without a history of cardiovascular disease residing in the northern part of the Netherlands^13^. All patients presented with a first ST-elevation myocardial infarction (STEMI) at the heart catheterization center at the University Medical Center Groningen. These patients are also part of the CardioLines Biobank^14^. Blood of these patients was collected in EDTA tubes (during hospital admission and 24 hours and 6−8 weeks after primary percutaneous intervention), and PBMCs were isolated using CPT tubes. PBMCs were cryopreserved until genotyping and scRNA-seq. Genotyping was done using the Infinium GSA (v1 for the samples processed with 10X 3’-end v2 chemistry and v3 for the samples processed with 10X v3 chemistry). scRNA-seq was conducted using 10X Genomics v2 (24 donors) and v3 chemistry (14 donors) reagents on sample pools of 7−8 donors each. Libraries were sequenced using a custom program (27-9-150 (v2), 28-8-150 (v3)) on an Illumina NovaSeq6000 using a 150-bp paired-end kit. Cell Ranger v3.0.2 software was used for basic alignment and initial processing of the scRNA-seq data. The processed (de-anonymized) scRNA-seq data, including a text file that links each cell barcode to its respective individual, has been deposited at the EGA under accession number [EGAD00001010064](https://ega-archive.org/datasets/EGAD00001010064). For the eQTL meta-analysis, only the scRNA-seq data from the 6−8 weeks timepoint was used, being most representative of healthy cells.

**Sawcer CSF cohort**

The Sawcer CSF cohort includes 81 adult patients with multiple sclerosis (33 patients) or other neurological diseases (48 patients) who were recruited from the Cambridge University Hospital (Addenbrooke’s) neurosciences department amongst those attending the department for a lumbar puncture as part of the investigation and treatment of their neurological condition^15^. Genotyping was done using the Infinium GSA. Cerebrospinal fluid was collected by lumbar puncture, and samples were either processed immediately or frozen for later pooling. To each sample, an equivalent volume of X-VIVO 10 serum-free hematopoietic cell medium was added to maintain cell viability. scRNA-seq was conducted using 10X Genomics 3’-end v2 chemistry reagents on sample pools of 1−4 donors each. Libraries were sequenced using a custom program (26-8-0-98) on an Illumina HiSeq4000 using a 75-bp paired-end kit. Cell Ranger v3.1 software was used for basic alignment and initial processing of the scRNA-seq data. The genotypes and raw scRNA-seq data have been deposited at the EGA under accession numbers [EGAD50000000183](https://ega-archive.org/datasets/EGAD50000000183) and [EGAD50000000184](https://ega-archive.org/datasets/EGAD50000000184), respectively. After QC (**Methods section: WP1-3**), 73 individuals remained for the eQTL meta-analysis.

**Franke multiome**

The Franke multiome dataset includes 128 adults from the general population of the northern part of the Netherlands, part of the Lifelines^16^ and Lifelines NEXT cohort^17^. Blood of these individuals was collected in EDTA tubes, and PBMCs were isolated using sodium-heparin CPT tubes. PBMCs were cryopreserved until genotyping and scRNA-seq. Genotyping owas performed using the Infinium GSA-MD v3. PBMCs, either left untreated or exposed for 24h to heat-killed *C. albicans blastoconidia*, were used for sc-multiomics analyses using 10X Genomics Multiome ATAC + Gene Expression v1 chemistry reagents on sample pools of 8 donors each. The single-nucleus RNA-seq (snRNA-seq) libraries were sequenced using a custom program (28-10-10-90) on a MGI-seq 2000 using a 100-bp paired-end kit. Cell Ranger arc 2.0.0 software was used for basic alignment and initial processing of the sc-multiomics data. CellBender v0.3.0 was used to correct the gene expression count matrix for systematic background noise. For this meta-analysis, only the snRNA-seq data from the untreated cells was used. After QC (**Methods section: WP1-3**), 118 individuals remained for the eQTL meta-analysis.

**Burkina Faso malaria cohort**

The Burkina Faso malaria cohort consists of 178 children from Burkina Faso recruited from rural

communities around Ouagadougou and Koudougou during the wet, high–malaria-transmission

season^18^. At the time of blood collection, all participants were tested for P.

falciparum infection. Peripheral blood was collected in heparin tubes, processed by Ficoll-Paque

density centrifugation, and cryopreserved in Burkina Faso before being shipped on dry ice to NYUAD for genotyping and scRNA-seq. Genomic DNA was genotyped using the H3Africa Infinium OmniExpress (24v13A1) array. scRNA-seq was performed on the 10x Genomics Chromium Controller using the Single Cell 3′ Library &amp; Gel Bead Kit v2, with 6 donors pooled per run, yielding a total of 32 pooled libraries. Sequencing was carried out on the Illumina NovaSeq 6000 using an S2 flow cell, and initial alignment and processing were performed using Cell Ranger v2.2. After QC **(Methods section: WP1-3)**, 178 individuals remained for the eQTL meta-analysis.
